# Dynamic extracellular proximal interaction profiling reveals Low-Density Lipoprotein Receptor as a new Epidermal Growth Factor signaling pathway component

**DOI:** 10.1101/2023.11.09.566449

**Authors:** Rasha Al Mismar, Payman Samavarchi-Tehrani, Brendon Seale, Vesal Kasmaeifar, Claire E. Martin, Anne-Claude Gingras

## Abstract

Plasma membrane proteins are critical mediators of cell-cell and cell-environment interactions, pivotal in intracellular signal transmission vital for cellular functionality. Proximity-dependent biotinylation approaches such as BioID combined with mass spectrometry have begun illuminating the landscape of proximal protein interactions within intracellular compartments. However, their deployment in studies of the extracellular environment remains scarce. Here, we present extracellular TurboID (ecTurboID), a method designed to profile cell surface interactions in living cells on short timescales. We first report on the careful optimization of experimental and data analysis strategies that enable the capture of extracellular protein interaction information. Leveraging the ecTurboID technique, we unveiled the proximal interactome of multiple plasma membrane proteins, notably the epidermal growth factor receptor (EGFR). This led to identifying the low-density lipoprotein receptor (LDLR) as a newfound extracellular protein associating with EGFR, contingent upon the presence of the EGF ligand. We showed that 15 minutes of EGF stimulation induced LDLR localization to the plasma membrane to associate with proteins involved in EGFR regulation. This modified proximity labelling methodology allows us to dynamically study the associations between plasma membrane proteins in the extracellular environment.

**One Sentence Summary:** We developed extracellular TurboID (ecTurboID) as a new proximity dependent biotinylation approach that can capture dynamic interactions at the cell surface, identifying Low-Density Lipoprotein Receptor as a new ligand-dependent extracellular partner of Epidermal Growth Factor Receptor.

## INTRODUCTION

The plasma membrane, enriched with a diverse array of proteins termed the ‘surfaceome’ (*1*), boasts receptors, enzymes, transporters, and adhesion molecules. These components are vital for cells to interface with their milieu, transmitting extracellular signals intracellularly. The dynamic and fluid nature of the plasma membrane allows cells to adapt to environmental changes (*2*) by interacting with the extracellular matrix (*3*, *4*), ligands (*5*) (e.g., cytokines and growth hormones), and neighboring cells (6). These interaction events are crucial in regulating signaling and driving cellular processes such as survival, proliferation, migration, or apoptosis (*1*). Yet, the dynamic nature of these plasma membrane protein interactions has posed considerable experimental challenges.

Historically, affinity purification (AP) combined with mass spectrometry (MS) emerged as a potent tool for protein complex analysis. However, its utility in studying cell surface interactions remains limited (7). Epitope tag-based AP-MS relies on fusing a protein of interest, known as a bait, to an epitope tag that is recognized by specific anti-epitope beads after cell lysis, typically using non-ionic detergents. Proteins complexing with the bait can then be captured and analyzed by MS. Due to the mild lysis conditions required to maintain the integrity of complexes through the AP step and subsequent washes, AP-MS is not well suited for resolving membrane protein complexes with a hydrophobic nature (7). Adaptation of AP-MS to study protein interactions at the plasma membrane would typically require stabilizing protein interactions, using crosslinking, for instance (*8*, *9*), which can elongate the process and convolute data interpretation, or require the systematic exploration of collections of detergents and lysis conditions that optimize recovery of specific complexes (*10*, *11*). Additionally, blue native gels have been used to identify membrane protein complexes (*12*, *13*) but require pre-solubilization of membrane proteins in mild lysis conditions to maintain complexes, presenting a similar limitation to affinity purification in studying proteins of hydrophobic nature.

Proximity-dependent biotinylation (PDB) techniques and MS offer broader coverage in studying cellular protein associations, including weak and membrane protein associations (*14*). PDB relies on fusing a modifying enzyme, such as a biotin ligase or a peroxidase, to a bait. These enzymes utilize biotin (biotin ligase) or biotin phenol (peroxidase) to induce covalent attachment of biotin-derived intermediate molecules to the bait and its adjacent proteins (preys) in living cells. Thus, harsh cellular lysis using ionic detergents or chaotropic agents can be used, with biotinylated proteins subsequently purified on streptavidin beads and identified by MS (*14*). This approach is ideally suited to solubilize and identify proximal interactors of plasma membrane proteins with hydrophobic nature (*15–18*).

The widely used PDB approach BioID employs an abortive *E. coli* biotin ligase (BirA R118G, BirA* (*19*)), with fast-acting derivatives miniTurbo and TurboID expanding the use of the approach for dynamic exploration (*20*). Abortive BirA enzymes convert biotin and adenosine triphosphate (ATP) to the reactive intermediate biotinyl-5’-AMP, which diffuses away from the enzyme’s active site and covalently binds to lysine residues on proteins within 10-20 nm of the bait (*21*). BioID has been used to identify new components of signaling pathways (*22*, *23*), uncover protein composition of membraneless organelles (*24*) and nuclear structures (*21*, *25*), identify proteins involved in lumen formation in 3D culture models (*26*), and map out the protein distribution in human cellular organelles (*18*). BioID was also applied to define the plasma membrane proteome by directing BirA* to the inner leaflet of the lipid bilayer (*16*) or by tagging the cytoplasmic portion of plasma membrane proteins to identify proteins in the vicinity of the membrane-spanning baits (*15*, *18*, *27*). However, these strategies generally overlook interactions occurring on the extracellular side of the plasma membrane, such as those involving the extracellular domain of membrane proteins, extracellular matrix proteins or proteins that are extracellularly tethered to the plasma membrane via a glycosylphosphatidylinositol (GPI) anchor.

Unlike the widespread and optimized use of BioID and other PDB techniques intracellularly, fewer but important studies have explored PDB extracellularly. PDB using horseradish peroxidase (HRP) coupled to a protein-specific antibody or a toxin has been used to identify molecular clusters at the cell surface (*28–30*) and profile lipid raft components (*31–33*). When coupled to MS, HRP-based PDB identifies protein interactions at synaptic clefts (*34*) and defines proteomes on the surface of neurons (*35*, *36*). The fact that biotin phenol has low membrane permeability and HRP is inactive in reducing environments such as the cytoplasm (*14*) makes HRP-based PDB applicable for cell surface profiling. While this approach has shown success in the extracellular space, HRP has a large labeling radius (200-300 nm) (*32*), making it better suited for defining surface proteomes (*35–37*) or large protein clusters (*28*, *29*) than specific protein associations. APEX2, an ascorbate peroxidase (*38*), induces labeling at the cell surface when tethered using lipidated DNA, yet to a much lower extent than HRP (*37*). Another approach to cell surface PDB is pupylation-based interaction tagging (PUP-IT) (*39*), which requires 24 hours of biotin treatment, limiting its application for dynamic interaction studies. TurboID, the fast-acting derivative of BirA* (catalyzes labeling in minutes compared to hours required by BirA*), was recently used to map ectodomain binding partners of E-cadherin (*40*) and to identify interacting proteins at astrocyte-neuronal junctions (*41*). BioID2, a biotin ligase from *Aquifex aeolicus* (*42*), also shows labeling potential when targeted to the extracellular side of the plasma membrane using a GPI anchor (*43*). These studies demonstrate the possibility of using BioID in extracellular space. However, the protocols used rely on long labeling times (hours’ time scale) – which is not applicable for dynamic studies – or use saturating biotin concentrations to achieve sufficient cell surface labeling – which may lead to high background signal (*20*).

In this context, we introduce extracellular TurboID (ecTurboID), a refined protocol that permits capturing dynamic extracellular protein partnerships within a short 15-minute labeling window. Harnessing the rapid action of TurboID biotin ligase, we profiled the extracellular proximal interactome of the tyrosine kinase receptor EGFR (epidermal growth factor receptor) under varying ligand conditions, unveiling LDLR (low-density lipoprotein receptor) as a novel EGF-dependent interactor. We further revealed that this interplay is orchestrated by an EGF-triggered shift in LDLR localization and its consequent association with other plasma membrane proteins.

## RESULTS

### Design and optimization of extracellular TurboID

To develop an extracellular biotin ligase PDB approach, we selected the TurboID-3xFLAG biotin ligase for its rapid labeling kinetics and efficiency. To position TurboID on the extracellular side of the plasma membrane, we fused it with an N-terminal signal sequence from immunoglobulin kappa (IgK). We appended a glycosylation motif (NNT: asparagine, glycine, threonine) to assist in sorting, posttranslational modifications, and membrane stabilization. As proof of principle, we cloned the transmembrane domain (TMD) from the bone morphogenic receptor type 1 A (BMPR1A, residues 149-179) in-frame with the ecTurboID, anchoring it to the plasma membrane. The resulting construct, ecTurboID-TMD (Fig. 1A, upper panel), was used to develop and optimize the ecTurboID approach.

**Figure 1:**
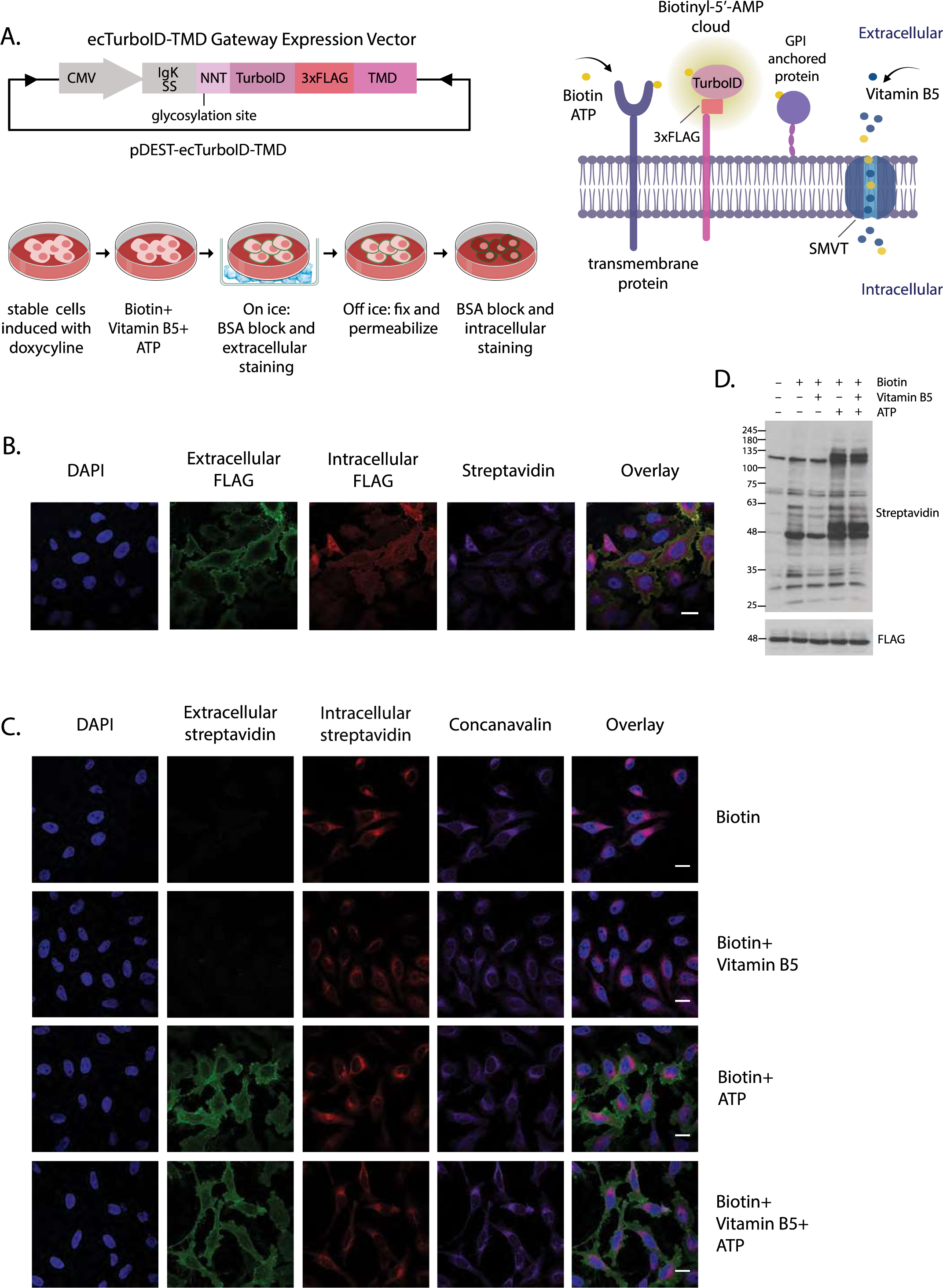
Vector design and optimization of extracellular TurboID in HeLa cells. (**A**) Top left: The transmembrane domain (TMD) of bone morphogenic receptor type 1 A (BMPR1A) was fused in frame with the ecTurboID (referred to as ecTurboID-TMD) under the control of doxycycline-inducible cytomegalovirus (CMV) promoter in a Gateway expression vector. NNT: asparagine, glycine, threonine. IgKSS: immunoglobulin kappa signal sequence. TurboID: biotin ligase. 3xFLAG: epitope tag. Top right: cartoon illustration of the labeling strategy for ecTurboID showing ecTurboID-TMD at the cell surface in the presence of ATP and vitamin B5. SMVT: sodium-dependent multivitamin transporter. Bottom left: cartoon illustration of the fluorescence protocol used in this study to show extracellular and intracellular expression and labeling of ecTurboID-TMD. **(B)** HeLa cells stably expressing ecTurboID-TMD were labeled for 30 minutes with 25 µM biotin and stained as illustrated in the bottom left cartoon in panel A, to show extracellular expression (green fluorescence) and intracellular expression (red fluorescence). Streptavidin was used to show intracellular labeling. Scale bar, 20 µm. **(C)** HeLa cells stably expressing ecTurboID-TMD were labeled with 25 µM biotin for 30 minutes in the presence of 5 mM vitamin B5 or 1.5 mM ATP, or both. Cells were stained with fluorescent streptavidin to show extracellular labeling (green fluorescence) and intracellular labeling (red fluorescence), as illustrated in the bottom left cartoon in panel A. Concanavalin was used as an ER marker. **(D)** Lysates from HeLa cells expressing TurboID-TMD labeled for 15 minutes with biotin in the presence of vitamin B5 and ATP (same concentrations stated in C) were probed for expression (FLAG) and biotinylation (HRP-conjugated streptavidin).

To distinguish between the extracellular and intracellular expression patterns of ecTurboID-TMD in Flp-In T-REx HeLa stable cell pools in which the addition of doxycycline-induced protein expression, we adapted a fluorescence technique to detect extracellular expression and biotinylated proteins in non-permeabilized cells, followed by membrane permeabilization and detection of intracellular expression and biotinylation (Fig. 1A, lower panel). We showed that ecTurboID-TMD is expressed at the cell surface and intracellularly (Fig. 1B), where it colocalized with intracellular streptavidin signal induced by the addition of 25 µM biotin for 15 minutes (Fig. 1B). PDB with biotin ligases requires ATP as a co-factor for the generation of the active biotinyl-5’AMP intermediate. Contrary to intracellular labeling that capitalizes on the cell’s intracellular ATP, ecTurboID requires exogenous ATP to catalyze labeling in the extracellular space. Here, we used 1.5 mM ATP to be near the range of intracellular ATP levels (range between 2 and 8 mM (*44*)); this concentration led to detectable extracellular biotinylation and was used for all experiments presented here.

In mammals, cellular biotin uptake is mediated by the sodium-dependent multivitamin transporter (SMVT; gene *SLC5A6*) (*45*), permitting intracellular proximity-dependent biotinylation. However, in cases where extracellular labeling is desired, biotin uptake can lead to intracellular labeling during the trafficking of the construct to the cell surface, which might mask the detection of cell-surface labeled proteins. SMVT is also a transporter of vitamin B5 (pantothenic acid) (*45*), and we reasoned that we could decrease biotin uptake and intracellular labeling by using vitamin B5 as a competitive inhibitor (Fig. 1A, upper right panel). To assess this strategy, we added increasing concentrations of vitamin B5 (millimolar concentration of vitamin B5 compared to micromolar concentrations for biotin) and monitored extracellular and intracellular labeling. Addition of vitamin B5 reduced intracellular labeling in a concentration-dependent manner (Fig. S1, A and B) while largely maintaining labeling at the cell surface (Fig. S2). The effect of increasing amounts of vitamin B5 was less pronounced beyond 25 µM biotin, likely indicating saturation in biotin levels. To ensure sufficient extracellular labeling while minimizing intracellular labeling, we used 25 µM biotin and 5 mM vitamin B5 throughout the study. To confirm the minimum labeling time required for sufficient extracellular signal visualization by fluorescence microscopy, we performed a time course of biotinylation that revealed robust extracellular labeling at 15 and 30 minutes following the addition of biotin and ATP, while intracellular labeling was more pronounced at 60 minutes, indicating that shorter labeling times would be preferable (Fig. S3).

Using streptavidin fluorescence microscopy, we illustrate the significance of adding biotin, vitamin B5 and ATP for 30 minutes to HeLa cells stably expressing ecTurboID-TMD (Fig. 1C). Intracellular labeling following biotin addition appeared in the ER and Golgi, as demonstrated by overlay with concanavalin. Biotin labeling is expected in the ER for proteins synthesized on ER-associated ribosomes and translocated in the ER for folding, glycosylation, and trafficking, which is the case for plasma membrane proteins. The addition of vitamin B5 decreased intracellular signal in the ER and Golgi, while adding ATP to the reaction induced labeling at the cell surface (Fig. 1C). In agreement with these findings, new biotinylated bands appear following the addition of ATP when characterized by far-western blot (Fig. 1D), while the addition of vitamin B5 reduced biotinylation levels, presumably through the decrease of intracellular labeling (Fig. 1D). In summary, using fluorescence microscopy, we developed an optimized protocol for extracellular labeling for PDB experiments.

### Mass spectrometric analysis of ecTurboID-TMD identifies cell surface proteins

We next used these optimized conditions to perform affinity purification and mass spectrometry analysis of ecTurboID-TMD-expressing cells treated with biotin. We first compared the enrichment of preys in the presence of ATP (enriching for extracellular labeling) to that in the absence of ATP in cells treated with vitamin B5 for 5, 15, 30 and 60 minutes through Gene Ontology (GO) Cellular Components analysis (Fig. S4A and B). While the recovery of plasma membrane and cell periphery terms was significant at each time point, the 15-minute time point demonstrated a more substantial enrichment of relevant terms (Fig. S4B). 524 preys were identified with ecTurboID-TMD from all conditions tested (biotin, biotin+vitamin B5, biotin+ATP, biotin+vitamin B5+ATP) at the 15-minute window. 152 identified proteins showed a ≥ 2-fold decrease in spectral counts (used as a proxy for abundance) following vitamin B5 addition, suggesting that they would be intracellularly labeled (Fig. 2A). In agreement with this, GO analysis of these depleted proteins retrieved components of the endomembrane system (Fig. 2D, first column). The addition of ATP led to the enrichment of 129 proteins, including those exclusively detected in the presence of ATP or that showed a ≥ 2-fold increase in spectral counts (Fig. 2B). ATP-dependent identifications were enriched for plasma membrane components, in addition to cell adhesion and receptor signaling processes, supporting the functionality of our approach at the plasma membrane and extracellular space (Fig. 2D, second column). While the addition of ATP identified a new set of proteins, the combination of ATP and vitamin B5 further bolstered their abundance, indicated by higher spectral counts for the ATP-dependent proteins (a subset of these preys is shown in Fig. 2E) and lower p-value for the enriched cellular components (Fig. 2D, third column), demonstrating that the decrease in intracellular labeling mediated by the addition of vitamin B5 improves the sensitivity of cell surface protein detection by MS (Fig. 2E). Cellular components showing ≥ 2-fold depletion in the presence of ATP alone or in combination with vitamin B5 further support the functionality of our system at the cell surface (Fig. S5). Collectively, we optimized the labeling conditions and duration that allowed the identification of cell surface and plasma membrane proteins by MS while minimizing intracellular protein labeling.

**Figure 2:**
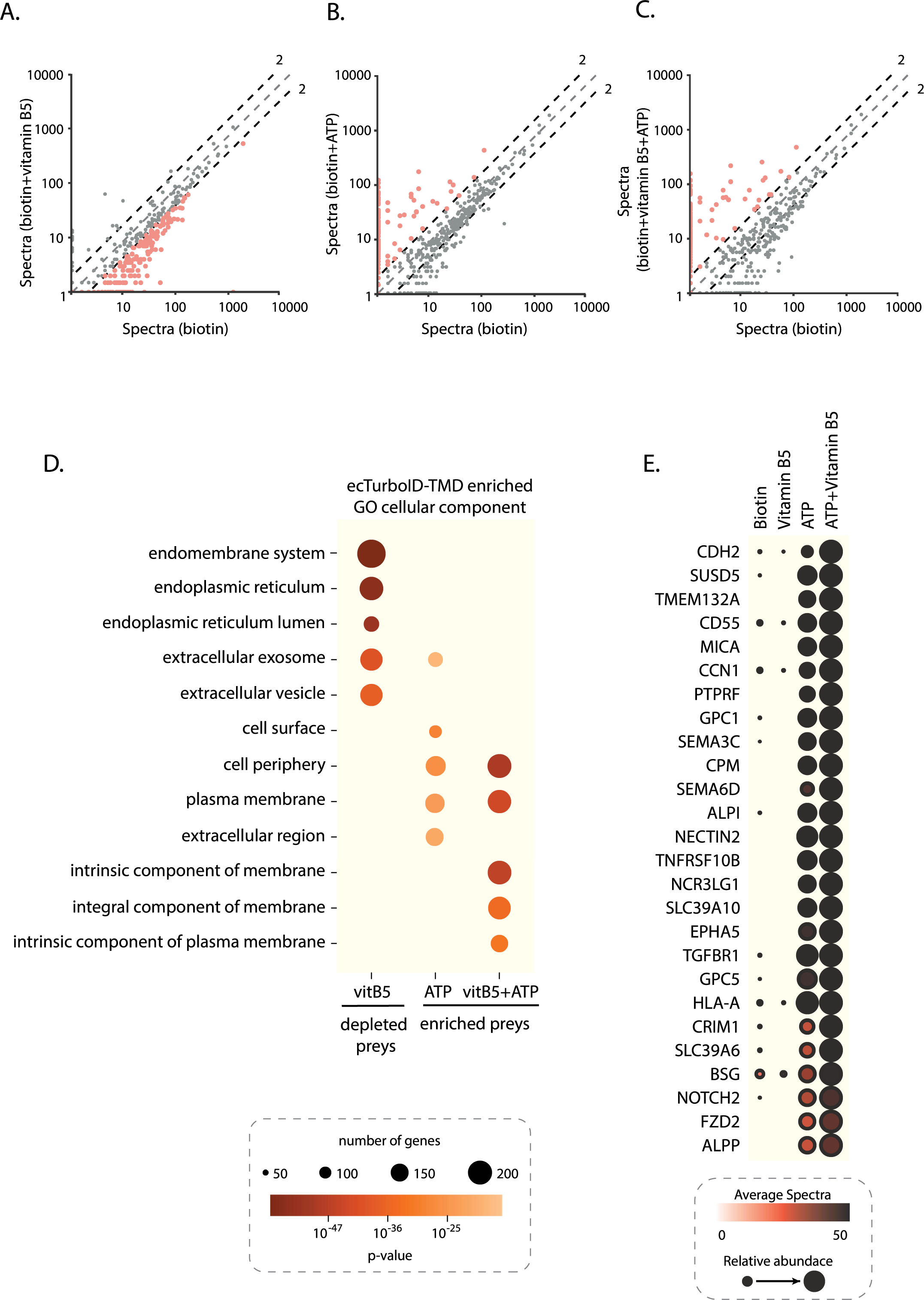
Identification of cell surface labeled proteins using TurboID-TMD by mass spectrometry. (**A–C**) Condition versus condition scatter plots (*68*) of the abundance of all proteins identified with 15 minutes of 25 µM biotin labeling in the presence of 5 mM vitamin B5 or 1.5 mM ATP (or both) in HeLa cells expressing ecTurboID-TMD, *n* = 2. The black dashed lines indicate the 2-fold change cutoff. The orange-colored dots indicate preys used for GO enrichment in D. (**D**) Gene Ontology (GO) analysis of cellular components enriched from preys with a 2-fold change or more upon addition of vitamin B5 (vitB5), ATP or both from (A–C). See the in-figure legend for the p-value and number of genes/proteins in each GO category. See Fig. S5 for additional GO enrichment analysis from this data. (**E**) Dot plot of a subset (26 out of 129) of the most abundant (by spectral counts) cell surface proteins identified in the presence of ATP, with and without vitamin B5 (same concentrations stated in D).

### Expressing TurboID to the plasma membrane with a GPI anchor labels cell surface proteins

We next assessed whether TurboID could also be anchored to the cell membrane via a GPI anchor by fusing the GPI anchoring signal sequence of complement decay accelerating factor (CD55) in frame with the engineered ecTurboID (ecTurboID-GPI; Fig. 3A). ecTurboID-GPI induced biotinylation at the cell surface in the presence of biotin and ATP (Fig. 3B, lower panel) and expression and biotinylation levels were comparable to that of ecTurboID-TMD (Fig. S6A). Using the SAINTexpress scoring method (*46*, *47*), 358 high-confidence proximity interactors were identified with either construct (Fig. 3C). Prey profiles identified with each construct in the absence of ATP (labeling in the endomembrane system) and those identified with eGFP (intracellular non-specific labeling) were used as controls for SAINT scoring. While gene ontology analysis of the high-confidence proximal interactors enriched for cellular components in the extracellular space (ecTurboID-GPI) and plasma membrane (ecTurboID-TMD), the enriched biological processes were distinct (Fig. 3D), suggesting that these constructs may localize to different domains of the plasma membrane. This hypothesis was further supported by preferential labeling of cell adhesion and GPI-anchored proteins by ecTurboID-GPI indicated by higher spectral counts (Fig. 3E).

**Figure 3:**
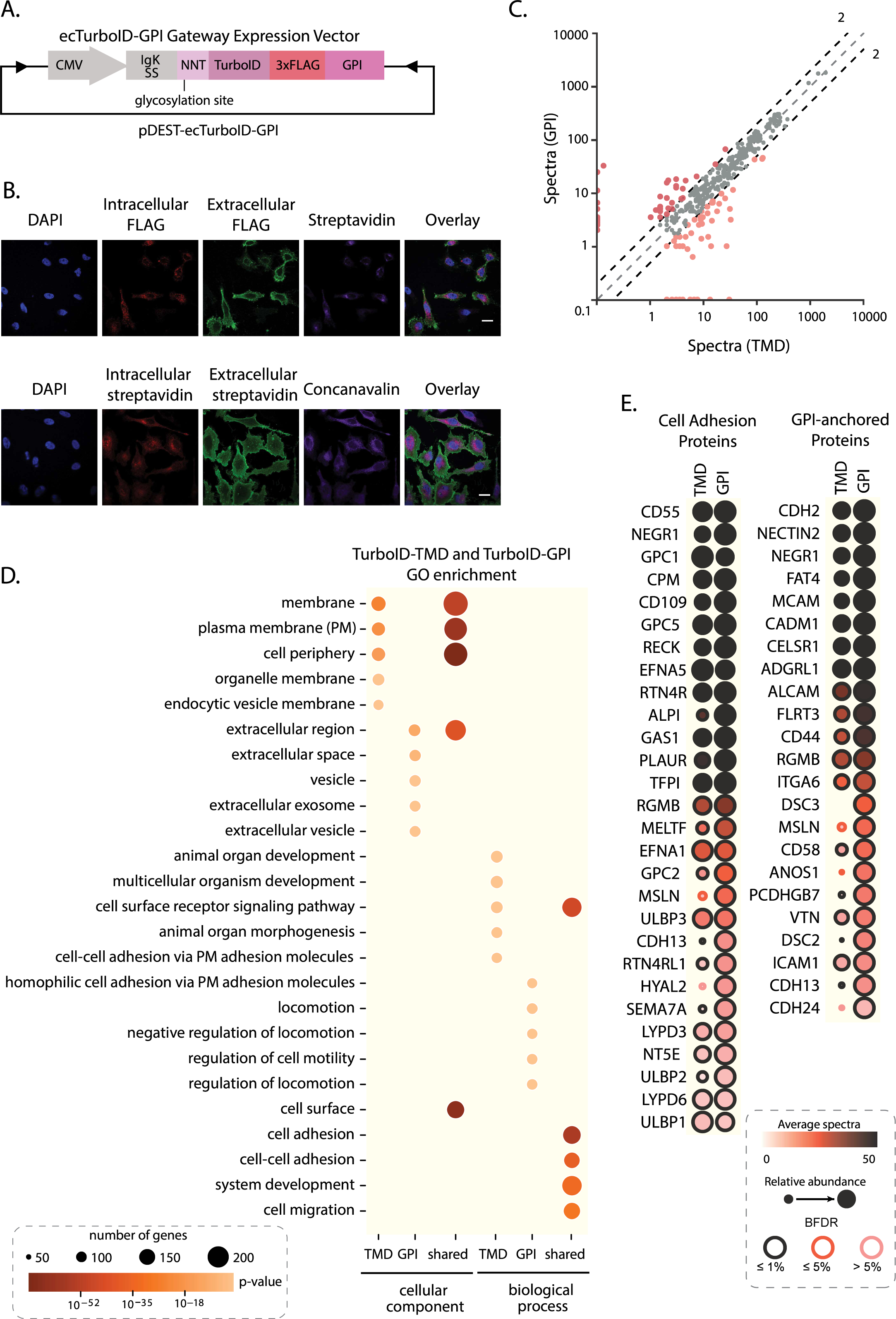
Anchoring ecTurboID at the plasma membrane using a glycosylphosphatidylinositol (GPI) anchor. (**A**) The GPI signal sequence from the complement decay accelerating factor was fused in frame with ecTurboID under the control of a doxycycline-inducible promoter in a Gateway expression vector. (**B**) Upper panel: HeLa cells stably expressing ecTurboID-GPI were labeled for 30 minutes with 25 µM biotin and stained to show extracellular expression (green fluorescence) and intracellular expression (red fluorescence). Fluorescent streptavidin was used to show intracellular labeling. Lower panel: HeLa cells stably expressing ecTurboID-GPI were labeled with biotin for 30 minutes in the presence of 5 mM ATP and stained to show extracellular labeling (green fluorescence) and intracellular labeling (red fluorescence) as illustrated in the bottom left cartoon in panel A of figure 1. Concanavalin was used as an ER marker. Scale bar, 20 µm. (**C**) Bait versus bait scatter plot of abundance (average spectral counts) for all high confidence proximity interactors (Bayesian False Discovery Rate, BFDR ≤ 1%, minimum of 2 spectral counts) identified with ecTurboID-TMD and ecTurboID-GPI after 15 minutes of 25 µM biotin, 5 mM vitamin B5 and 1.5 mM ATP, *n* =2. The black dashed lines indicate the 2-fold change cutoff. The colored dots indicate preys used for GO enrichment in D. (**D**) Gene Ontology (GO) analysis for cellular compartments and biological processes enriched from high-confidence proximity interactors (BFDR ≤ 1% and ≥ 2-fold enriched) identified with ecTurboID-TMD, ecTurboID-GPI and both (shared) after 15 minutes of biotin labeling in the presence of vitamin B5 and ATP in HeLa cells (same concentrations stated in C). (**E**) Dot plot (columns: baits, rows: proximal interactors) of a manually curated list of GPI-anchored and cell adhesion proteins identified with either construct (BFDR ≤ 1% and a minimum of 2 spectral counts) when cells were labeled with biotin for 15 minutes in the presence of vitamin B5 and ATP (same concentrations stated in C).

### ecTurboID can be applied to extracellularly profile different type 1 membrane receptors

To explore the specificity of ecTurboID, we extracellularly tagged and profiled five type 1 membrane proteins that are expressed in HeLa cells (*48*): the receptor tyrosine kinases epidermal growth factor receptor 1 (EGFR) and epidermal growth factor receptor 2 (HER2 or ERBB2), the serine/threonine kinase bone morphogenic receptor type 1A (BMPR1A), the multifunctional non-receptor transmembrane protein Basigin (BSG or CD147), and the low-density lipoprotein receptor (LDLR). All proteins could localize to the cell surface and induce biotinylation at the plasma membrane (Fig. 4A and Fig. S6B). However, BMPR1A was expressed at a low level, leading to a weaker biotinylation signal than the other baits. For SAINT scoring, we used prey profiles identified with each bait of interest in the absence of ATP and those identified with eGFP as controls. In total, we identified 214 high-confidence proximity interactors across all baits. Gene ontology analysis of the high confidence interactions enriched for cellular components, biological processes, and molecular functions specific to the plasma membrane (Fig. 4B). While many preys were shared across all baits (Fig. 4C), unique associations were observed, including the identification of apolipoproteins with LDLR, BMPR1B with BMPR1A, and EGFR with ERBB2 (Fig. 4C and Fig. S6C). We further compared the proximal interactomes of EGFR, ERBB2 and LDLR, which shared similar expression levels (Fig. S6B). EGFR and ERBB2 showed comparable proximal interactomes, while LDLR held a distinct profile showing apolipoproteins as specific proximity interactors (Fig. 4C and D). Together, these experiments reveal that our ecTurboID approach is applicable to type I transmembrane proteins and yields meaningful proximal interactome data.

**Figure 4:**
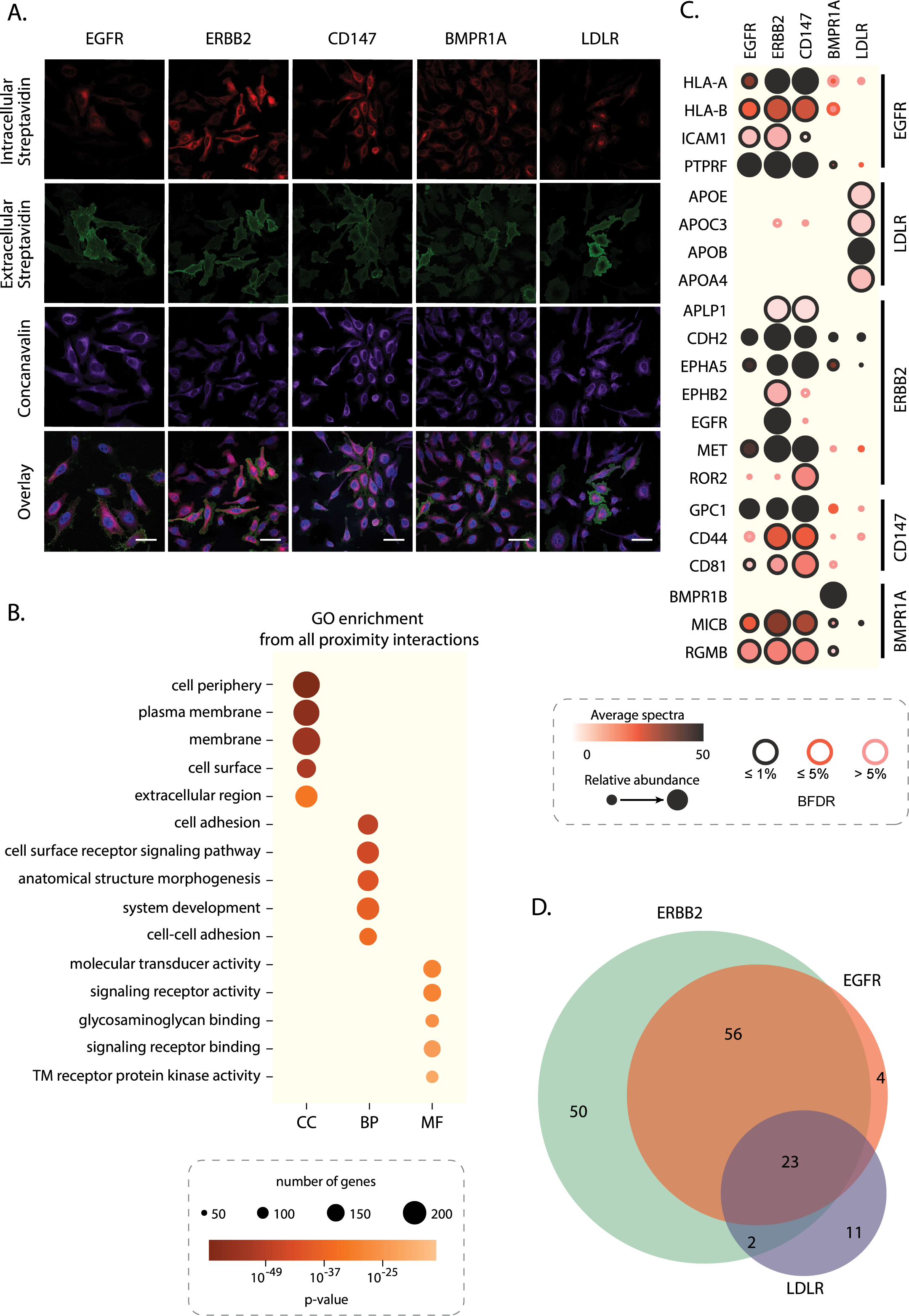
Profiling type 1 membrane receptors using ecTurboID. (**A**) HeLa cells stably expressing EGFR, ERBB2, CD147, BMPR1A or LDLR fused in frame with ecTurboID were labeled with 25 µM biotin for 30 minutes in the presence of 1.5 mM ATP and stained to show extracellular labeling (green fluorescence) and intracellular labeling (red fluorescence), as illustrated in the bottom left cartoon in panel A of figure 1. Concanavalin was used as an ER marker. Scaler bar, 20µm for EGFR and 50 µm for other baits. (**B**) Gene Ontology (GO) analysis of cellular components (CC), biological processes (BP) and molecular functions (MF) enriched from high confidence proximity interactors (BFDR ≤ 1% and a minimum of 2 spectral counts) identified with all baits from HeLa cells treated with 25 µM biotin, 5 mM vitamin B5 and 1.5 mM ATP for 15 minutes. TM: transmembrane. (**C**) Dot plot (columns: baits, rows: proximal interactors) of a subset of previously known interactors (for each bait, shown on the right) identified as high confidence (BFDR ≤ 1% and minimum of 2 spectral counts) after 15 minutes of biotin, ATP, and vitamin B5 (same concentrations stated in B), *n* = 2. (**D**) Venn diagram of all high confidence proximity interactors (BFDR ≤ 1% and minimum of 2 spectral counts) identified with EGFR, ERBB2 and LDLR after 15 minutes of biotin, ATP, and vitamin B5 (same concentrations stated in B).

### ecTurboID of EGFR reveals lipid homeostasis proteins as high-confidence interactions upon EGF stimulation

We then tested the functionality of ecTurboID in detecting dynamic interactions at the cell surface by profiling EGFR in the steady state and following stimulation by its ligand EGF. To complete the EGFR interaction profile and highlight new extracellular interactions identified by ecTurboID, we also profiled it intracellularly (we refer to extracellularly-tagged EGFR as EGFR-N since we tagged its N-terminus, and the intracellularly-tagged as EGFR-C; Fig. 5A). For intracellular profiling of EGFR, we used TurboID and the natural N-terminus of EGFR. Like EGFR-N, EGFR-C localized to the plasma membrane and induced labeling upon addition of biotin (Fig. 5B-C). We used both a low (0.25 ng/ml) and a high (10 ng/ml) concentration of EGF for stimulation since receptor activation, clustering, and internalization are dependent on the dose of EGF used (*49*). EGFR tagged at the N or C terminus was able to bind EGF and become phosphorylated at tyrosine 1068, a marker of activated EGFR (Fig. 5D). We identified a total of 602 high-confidence proximity interactors with either EGFR-N or C across conditions (93 with EGFR-N, 468 with EGFR-C and 41 with both). For SAINT scoring, we used prey profiles identified with EGFR-N in the absence of ATP and those identified with eGFP as controls. Distinct interaction profiles were identified with EGFR on each side of the plasma membrane (Fig. 6A). Cell adhesion and cell surface receptor signaling were among the top enriched biological processes from EGFR-N (extracellular) proximity interactions while signaling and cytoskeleton organization processes were enriched in interactors specific to EGFR-C (Fig. 6B). Proximity interactions identified by both EGFR-N and C enriched for biological processes at the extracellular and intracellular space, respectively, reflecting the transmembrane nature of these proteins; 53 out of 59 commonly identified proximity proteins have both an extracellular domain and a cytoplasmic tail, allowing their labeling on either side of the plasma membrane.

**Figure 5:**
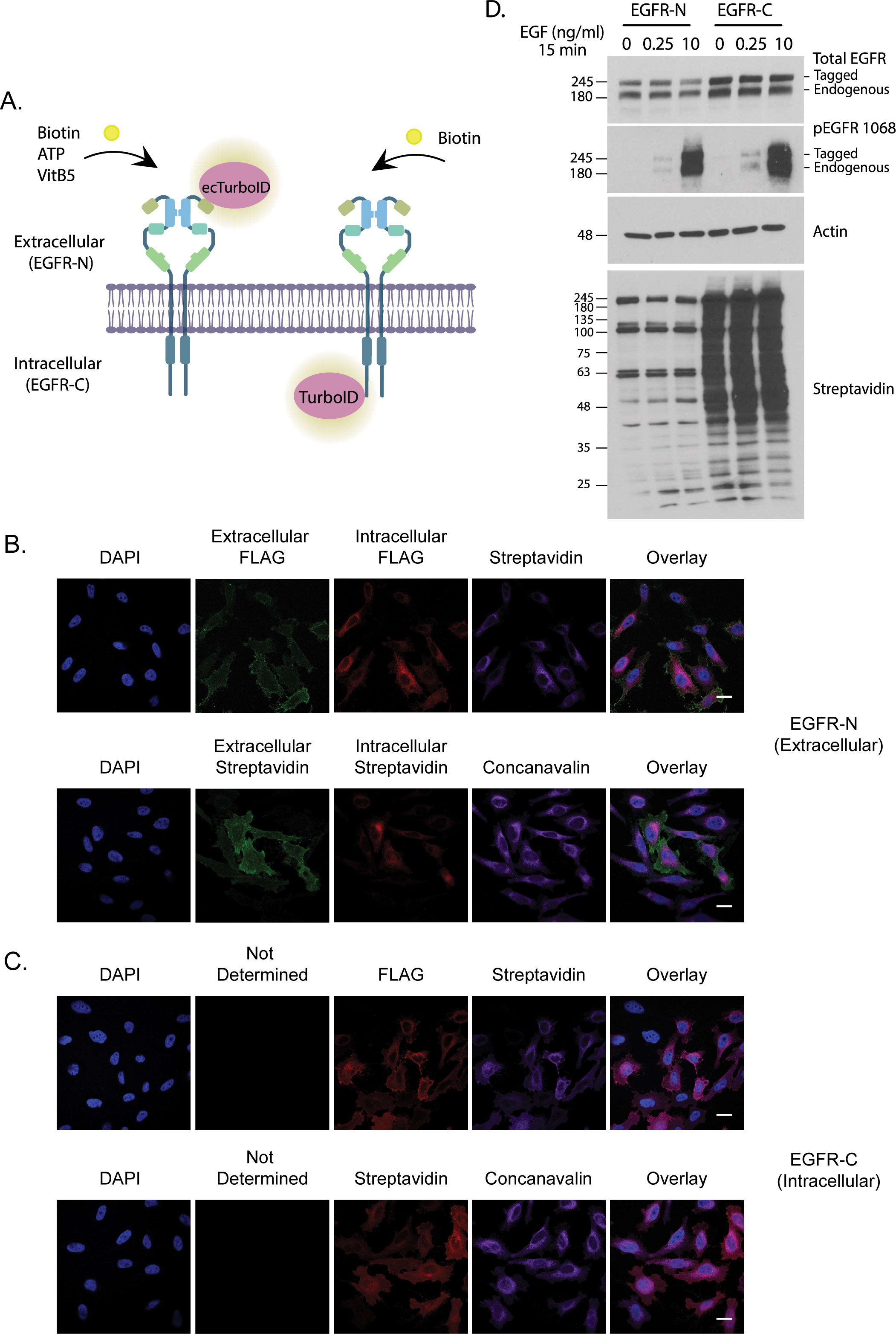
Expression, labeling and EGF stimulation of tagged EGFR. (**A**) Cartoon illustration of EGFR at the plasma membrane showing tagging at both termini and the labeling conditions used for each. (**B**) Upper panel: HeLa cells stably expressing N-tagged EGFR were labeled with 25 µM biotin and stained for extracellular expression (green fluorescence) and intracellular expression (red fluorescence). Streptavidin was used to show intracellular labeling. Lower panel: cells labeled with biotin in the presence of 1.5 mM ATP were stained to show extracellular (green fluorescence) and intracellular labeling (red fluorescence), as illustrated in the bottom left cartoon in panel A of figure 1. Concanavalin was used as an ER marker. Scale bar, 20µm. (**C**) HeLa cells expressing C-tagged EGFR were labeled with 25 µM biotin and stained to show expression and labeling (red fluorescence). Scale bar, 20µm. (**D**) HeLa cells stably expressing N-tagged or C-tagged EGFR were labeled with 25 µM biotin and EGF stimulated for 15 minutes in the presence of 1.5 mM ATP (for N-tagged). Cell lysates were used to probe for EGFR stimulation and biotin labeling.

**Figure 6:**
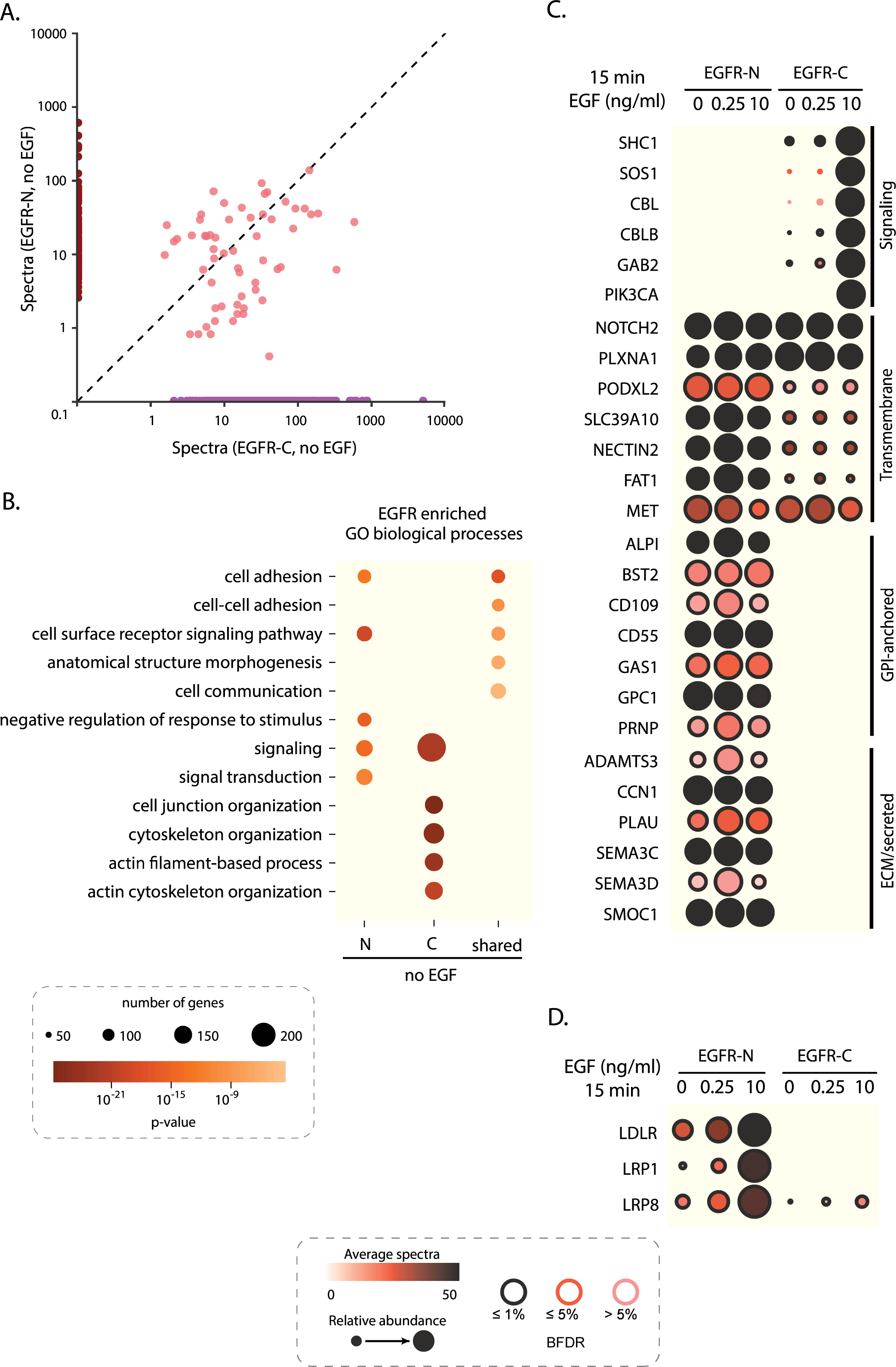
Profiling of ecTurboID-tagged EGFR in response to EGF stimulation. (**A**) Bait versus bait scatter plot of abundance (average spectral counts) for all high confidence proximity interactors (BFDR≤ 1% and a minimum of 2 spectral counts) identified with EGFR-N and EGFR-C after 15 minutes of 25 µM biotin (in addition to 5 mM of vitamin B5 and 1.5 mM ATP for N-tagged expressing HeLa cells) without stimulation. (**B**) Gene Ontology (GO) analysis of biological processes enriched from high confidence proximity interactors shown in panel A. (**C**) Dot plot (columns: baits, rows: proximal interactors) of a subset of high confidence proximal interactors (manually curated from each category) identified with both tagged proteins from cells labeled with biotin (in addition to vitamin B5 and ATP for EGFR-N expressing HeLa cells, same concentrations stated in A) for 15 minutes in the presence or absence of EGF. (**D**) Dot plot of lipid metabolism proteins identified with EGFR-N high confidence proximal interactors, *n* = 2.

Proteins involved in EGFR signaling and ubiquitination were detected with EGFR-C in a ligand-dependent manner (Fig. 6C), as expected. GPI-anchored proteins, extracellular matrix components and secreted proteins were only recovered in the EGFR-N ecTurboID. Proteins identified extracellularly showed a higher association with EGFR, reflected by higher spectral counts, after treatment with 0.25 ng/ml EGF, likely due to receptor clustering, in comparison to 10 ng/ml EGF, which is known to induce EGFR internalization (*49*) and lead to loss of association with plasma membrane proteins (Fig. 6C). On the other hand, proteins involved in cellular lipid metabolism, including LDLR, low-density lipoprotein receptor-related protein 1 (LRP1) and low-density lipoprotein receptor-related protein 8 (LRP8), associated with EGFR in an EGF dose-dependent manner, with maximal association at 10 ng/ml EGF (Fig. 6D). LRP1 and LRP8, but not LDLR, have been previously reported as interactors of EGFR (*50*). However, a functional association between EGFR and LDLR has been established in breast cancer and glioblastoma, where prolonged EGFR stimulation increases total LDLR expression (*51*, *52*). For these reasons, we further characterized the relationship between LDLR and EGFR.

### EGF induces re-localization of LDLR after 15 minutes of stimulation

To assess whether the increased proximal interaction detected for EGFR and LDLR could simply reflect modulated LDLR levels, we first confirmed that our short 15 minutes of EGF stimulation did not increase total LDLR expression (Fig. S7A). Next, after EGF stimulation, we performed reciprocal TurboID analysis of LDLR, extracellularly or intracellularly tagged. We used 20 ng/ml EGF to improve potential associations of LDLR and EGFR, as this association was EGF dose-dependent. EGF stimulation did not affect proximal interactions detected with the extracellularly-tagged LDLR (Fig. S7B). On the other hand, a total of 623 high-confidence proximity interactors (including EGFR) were identified with LDLR-C (intracellular) in either the resting or stimulated condition (Fig. 7A). We used prey profiles identified with LDLR-N in the absence of ATP and those identified with eGFP as controls for SAINT scoring. Contrary to results for EGFR-N, EGFR’s identification as a prey by LDLR-C did not change following the addition of EGF; however, many other proteins showed regulated associations (Fig. 7B). Among the high confidence proximity interactors, 35 showed a ≥ 2-fold increase in spectral counts upon EGF stimulation while 27 showed a ≥ 2-fold decrease (Fig. 7A). EGF-dependent increase in proximity interactions enriched for components of the plasma membrane, indicating a change in LDLR localization (Fig. 7C). Proteins involved in EGFR signaling, cytoskeleton organization and phosphoinositide kinase (PiK) showed increased association with LDLR after EGF stimulation. In addition, dedicator of cytokinesis (DOCK) proteins, which act as guanine exchange factors (GEF) to activate Rho GTPases, also associated with LDLR upon EGF stimulation (Fig. 7B). In contrast, members of the exocyst complex (EXOC3, EXOC4, EXOC6 and EXOC6B), which docks proteins to the plasma membrane, and Rab GTPase activation proteins showed reduced association with LDLR after EGF stimulation (Fig. 7B), further indicating a change in LDLR localization. When comparing the intracellular interactome of EGFR and LDLR, 40% of preys identified with LDLR were shared with EGFR in resting and EGF-stimulation conditions (Fig. 7D). We identified a subset of proteins which showed a ≥ 2-fold EGF-dependent increase in spectral counts with both EGFR and LDLR (Fig. 7D), including Girdin (CCDC88A), ERBB receptor feedback inhibitor 1 (ERRFI1) and mitogen-activated protein kinase kinase kinase kinase 5 (MAP4K5), all of which are previously known EGFR interactors (*50*). ERRFI1 is a negative regulator of EGFR signaling (*53*) while CCDC88A is a non-receptor guanine nucleotide exchange factor that localizes to the plasma membrane upon EGF stimulation (*54*) and complexes with G(i) alpha subunit and EGFR to promote cell migration (*55*). We further confirmed by immunofluorescence that EGFR and LDLR colocalize after 15 minutes of EGF stimulation (Fig. S7C). Altogether, these findings support the use of extracellular TurboID to identify new signaling components that might have been overlooked with currently available techniques.

**Figure 7:**
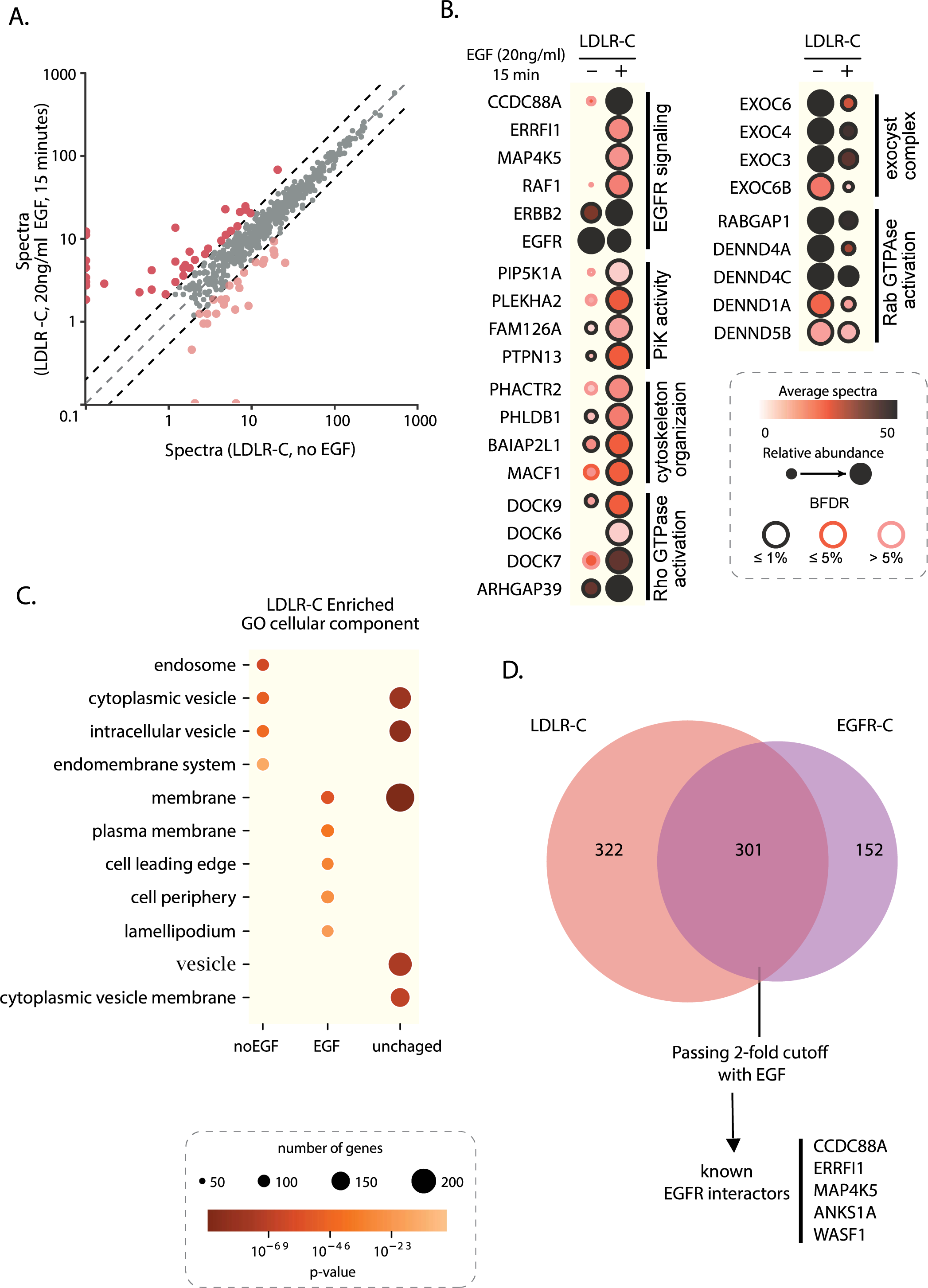
Profiling LDLR with and without EGF stimulation. (**A**) Condition versus condition scatter plot of abundance (average spectral counts) for all high confidence proximity interactors (BFDR ≤ 1% and a minimum of 2 spectral counts) identified after 15 minutes of 25 µM biotin in LDLR-C expressing HeLa cells with and without EGF stimulation, *n* = 3. The black dashed lines indicate the 2-fold change cutoff. The colored dots indicate preys used for GO enrichment in C. (**B**) Dot plot (columns: baits, rows: proximal interactors) of a subset of high-confidence proximal interactors identified after 15 minutes of 24 µM biotin in the presence or absence of 20 ng/ml EGF stimulation in LDLR-C expressing HeLa cells. (**C**) Gene ontology (GO) analysis of cellular components enriched from high confidence proximity interactors (BFDR ≤ 1%) after 15 minutes of 25 µM biotin in the presence or absence of EGF in LDLR-C expressing HeLa cells. (**D**) Venn diagram of all high-confidence proximity interactors identified with EGFR-C and LDLR-C expressing HeLa cells labeled for 15 minutes with 25 µM biotin in the resting and EGF stimulated conditions highlighting a subset of proteins that show a ≥ 2-fold increase in spectral counts with both tagged proteins upon EGF stimulation.

## DISCUSSION

Over the past decade, PDB coupled with MS has become a powerful tool for uncovering protein interactions underlying cellular processes and identifying organellar protein compositions (*56*). PDB-MS has been used to explore global protein changes and associations at the cell surface (*35–37*, *41*). However, its application to study dynamic cell surface protein interactions is still not widely explored (*14*). To complement currently available toolkits for cell surface profiling, we have now added ecTurboID. This powerful strategy allows the profiling of dynamic proximal protein interactions at the extracellular side of the plasma membrane using the fast-acting enzyme TurboID within short timescales (*20*). During the development of ecTurboID, two published studies demonstrated the feasibility of using TurboID at the cell surface to profile N-cadherin ectodomain partners and proteomes at neuronal junctions (*40*, *41*). Our approach, however, differs in its versatile application to single-pass type 1 and GPI-anchored proteins, using the accessible and efficient Gateway cloning system. ecTurboID also offers robust cell surface biotinylation within minutes of labeling compared to hours used before, permitting the study of dynamic interactions at the cell surface. We highlight ATP and vitamin B5 as critical components required for biotinylation in the extracellular space and minimizing intracellular labeling, respectively. Finally, we implemented a statistical scoring approach to enrich high-confidence interactions at the cell surface.

To catalyze labeling at the cell surface, adding ATP was essential to inducing biotinylation. The lack of cell surface labeling in the absence of ATP helped define controls for probabilistic scoring using SAINTexpress (*47*). Combined with vitamin B5 treatment, this approach resulted in enriched interaction profiles at the cell surface. To further validate our approach, fusing ecTurboID with a GPI-anchor proved that ecTurboID could be extended to study a set of proteins often overlooked in the currently available BioID-MS approaches targeting the plasma membrane. GPI-anchored proteins are known to cluster into cholesterol-rich regions known as lipid rafts and play a role in cell adhesion and signaling (*57*). The enrichment of GPI-anchored and cell adhesion proteins with ecTurboID-GPI compared to ecTurboID-TMD suggests that these constructs localize to different areas on the plasma membrane, highlighting a more targeted application of ecTurboID.

One limitation we faced with ecTurboID, like other tagging systems, is the difference in expression levels between the tagged proteins, affecting the labelling efficiency. For instance, CD147 showed the highest level of expression and thus led to a higher recovery of proteins that were shared with the other baits profiled in our study. BMPR1A had the lowest expression level and lowest number of identifications, yet it was the only bait to label BMPR1B, indicating specificity. LDLR exhibited a unique interaction profile compared to EGFR and ERBB2, both of which shared a similar profile consistent with the potential for heterodimerization in this receptor tyrosine kinases family. Therefore, the specificity of ecTurboID profiles should be determined by comparing proteins with similar and distinct functions at comparable expression levels (e.g. through the use of regulatable expression systems).

Profiling EGFR with and without EGF demonstrated the applicability of ecTurboID in determining dynamic interaction profiles in short timescales. Peroxidase-based PDB also catalyzes labeling in minutes but utilizes hydrogen peroxide to create biotin phenol radicals, which may be toxic in living cells and modulates signal transduction events. In addition, covalent tagging of tyrosine residues by biotin phenol radicals makes the use of peroxidases less desirable in PDB studies involving posttranslational modifications (*56*). We saw a distinct interaction profile of EGFR on the cell surface versus intracellularly. LDLR was among other lipid metabolism proteins that were identified with EGFR extracellularly in an EGF-dependent manner. This association has not been reported until recently. EGFR and LDLR are identified as interacting proteins on the surface of pancreatic adenocarcinoma cell lines using interaction-guided cross-linking coupled to AP-MS (*9*). It is worth noting that LDLR has a short cytoplasmic tail (50 amino acids) with 3 lysine residues. Given that biotin covalently attaches to lysine residues on vicinal proteins, LDLR might be missed in methods targeting the cytoplasmic side of the plasma membrane, as shown here with cytoplasmically-tagged EGFR, highlighting the importance of approaches such as ecTurboID in identifying new interactions.

Previous work demonstrates that LDLR expression can be upregulated by EGF-EGFR signaling in breast cancer via the MAPK pathway (*52*) and in glioblastoma and lung cancer via EGFR-associated PI3K/Akt signaling (*51*, *58*). While we showed that 15 minutes of EGF stimulation did not affect total LDLR expression, the change in LDLR interactome indicated a change in its localization. LDLR lost association with exocyst complex proteins and gained proximal interactions with proteins involved in EGFR signaling and cytoskeleton organization proteins. The association and colocalization we saw between EGFR and LDLR after EGF stimulation indicate that the two proteins may reside in compartments near or at the plasma membrane, within the time frame of stimulation we used in this study. The fact that we did not use saturating concentrations of EGF in our stimulation experiments may have resulted in unliganded EGFR being retained at the plasma membrane, where it associated with LDLR after EGF stimulation. Contrary to this observation, we did not see an EGF-dependent association between EGFR and LDLR when we profiled the latter with TurboID, possibly due to faster EGFR internalization at the higher EGF concentration used for reciprocal TurboID. Additionally, the intracellular interactome of LDLR reflects interactions from the total cellular pool of LDLR, which might dilute interactions at the cell surface. Our data and previously published reports suggest a role for LDLR beyond cholesterol delivery into the cell. Consistent with this hypothesis, LRP-1, another LDLR family member that was also identified as an EGF-dependent EGFR proximal interactor in our dataset, has been recently shown to stabilize activated EGFR at the plasma membrane which leads to activation of pro-motility signaling (*59*).

We optimized and validated ecTurboID coupled to MS as a new approach to profile proximal interactions in the extracellular space. ecTurboID is applicable to studying a broad range of proteins at the plasma membrane, and we anticipate that it will be valuable for investigating extracellular protein associations of type 1 membrane and GPI-anchored proteins and their role in signaling. Given the dynamic spatiotemporal organization of plasma membrane proteins, which might affect the specificity of interactions identified for each protein, ecTurboID is best suited to profile and compare a large cohort of plasma membrane proteins or study interactions contextually and dynamically, such as ligand stimulation, drug treatment or genetic depletion. In addition, ecTurboID could be adapted to profile intercellular (trans) interactions between cells, compared to cis interactions on the same cell, using appropriate labeling and quantitative proteomic approaches.

## MATERIALS AND METHODS

### ecTurboID vector design and Gateway cloning

pcDNA5-pDEST Gateway vector harboring TurboID-3xFLAG sequence under the control of a doxycycline-inducible promoter was used to design the ecTurboID destination vector. TurboID-specific primers were designed such that the forward primer included NheI restriction sites, no ATG, and a glycosylation site (NNT), while the reverse primer included an AscI restriction site. The PCR-amplified product was ligated into pcDNA5-pDEST digested with NheI and AscI. Immunoglobulin kappa (IgK) signal sequence (SS) was then added upstream of the modified TurboID using restriction digestion. Briefly, two complementary oligos with the IgK signal sequence, a Kozak consensus sequence, and KpnI and NheI restriction sites were annealed and ligated into the destination vector previously cut with the same enzymes. The resulting vector, ecTurboID, was used to make expression vectors for all extracellularly tagged baits. To design TurboID-TMD, TMD was amplified from the bone morphogenic receptor type 1A (BMPR1A) sequence-verified entry clone (NM_004329.3) by PCR using sequence-specific primers flanked by *att*B sites and a stop codon in the reverse primer. The PCR product was then cloned into a pDONR223 vector to create a new entry clone. Using LR Gateway cloning, TMD was transferred to the ecTurboID destination vector. Similarly, tagged proteins generated by Gateway cloning from sequence validated entry vectors generated by PCR from cDNA clones, EGFR (BAI46646.1), ERBB2 (NM_004448.4), CD147 (Basigin) (NM_198598.3), LDLR (NM_000527.5) and BMPR1A (NM_004329.3) were fused in frame with the ecTurboID construct. Sequence-specific forward primers with *att*B sites were designed downstream of the signal sequence, and the reverse primer had a stop codon. The PCR product was then cloned into pDONR223 and then into the ecTurboID destination vector. To fuse ecTurboID with a GPI anchor, a gene fragment with sequences specific to the GPI attachment signal from complement decay accelerating factor (CD55) flanked by *att*B sites was cloned into pDONR223 then into the ecTurboID destination vector. Enhanced green fluorescence protein (eGFP) and cytoplasmically-tagged EGFR and LDLR expression vectors were made by direct cloning from entry clones into pcDNA5-pDEST-TurboID. All primers and oligo sequences are shown in Table S1. Whole plasmid sequencing was performed on all expression vectors at Plasmidsaurus (https://www.plasmidsaurus.com/).

### Cell Culture and labeling conditions

Flp-In-T-REx HeLa cells were grown in complete growth media consisting of DMEM with 4.5 g/L glucose and 4 mM L-glutamine (Thermo Scientific, Cat# 11965) supplemented with 5 % fetal bovine serum (FBS), 5 % Cosmic calf serum (CCS), 100 U/mL penicillin and 100 µg/mL streptomycin. The Flp-In and T-REx loci were maintained in media containing 100 µg/mL zeocin (Life Technologies, Cat# R25001) and 3 µg/mL blasticidin (Bioshop, Cat# BLA477.100) prior to establishing stable lines. Cells transfected with Gateway expression vectors using jetPRIME Transfection Reagent (Polyplus Transfection) were selected and maintained in media containing 200ug/ml hygromycin B. For bait expression induction, cells were incubated in biotin depleted media containing 1 µg/ml doxycycline for 24 hours prior to biotin labeling. Cells were then labeled with biotin (BioBasic, Amherst, New York, Cat# 58-85-5) in the presence or absence of ATP and vitamin B5. Flp-In-T-REx HeLa cells stably expressing EGFR or LDLR were induced with doxycycline for 24 hours then serum starved overnight prior to stimulation with EGF (Sigma Aldrich, Cat# E9644) for the same time of biotin labeling.

### Serum biotin depletion

Fetal bovine serum and cosmic calf serum were mixed in a 1:1 ratio and incubated with pre-washed Streptavidin agarose beads (GE Healthcare Life Science, Cat# 17511301). Briefly, the required amount of beads was made into a 50% slurry and washed 3 times with sterile phosphate buffered saline (PBS). 10 µl of streptavidin bead slurry was used to deplete 5 ml of serum by incubating for 3 hours or overnight with gentle shaking. The serum was then sterile filtered and used to make biotin depleted media.

### Fluorescence staining of live and fixed cells

A fluorescence staining protocol was adapted to stain cells both extracellularly and intracellularly. Briefly, Flp-In-T-REx HeLa cells were plated on glass coverslips and the next day expression was induced with doxycycline for 24 hours then treated with biotin, ATP, and vitamin B5 for 30 min. Cells were washed twice with ice cold phosphate buffered saline (PBS) containing 200 mM calcium chloride and 100 mM magnesium chloride (PBS +/+) on ice. Cells were then blocked with 1.5 % bovine serum albumin (BSA) in PBS +/+ for 10 min on ice and stained with Alexa Fluor 488 conjugated streptavidin (Molecular Probes, ThermoFisher Scientific, A11001, 1:1000). For bait expression, mouse anti-FLAG (Monoclonal anti-FLAG M2 antibody, Sigma-Aldrich, Cat# F3165, 1:750) was added in blocking solution for 15 minutes followed by washing with 0.05% BSA in PBS +/+, then incubated with Alexa Fluor 488 secondary anti-mouse (Invitrogen, Cat# A11001, 1:1000) for 10 minutes. Cells were then fixed with 4 % PFA in 1X PBS +/+ for 20 minutes and permeabilized with 0.2 % NP-40 for 10 minutes. For intracellular streptavidin staining, cells were blocked for 30 min in 2 % BSA in PBS then stained with Alexa Fluor 594 conjugated streptavidin (Invitrogen, Cat# S11227, 1:2500), Alexa Fluor 647 conjugated concanavalin A (Thermo Fisher, Cat# C21421, 1:500) and DAPI (Sigma Aldrich, 20 mg/ml, used at 1:10000) in blocking solution for 1 hour. For intracellular FLAG staining, cells were blocked with 2% milk in PBS for 30 minutes. Mouse anti-FLAG (1:2000) was added in blocking solution for 3 hours rotating on ice. Cells were then washed 3X with 0.05% BSA in PBS and stained with Alexa Fluor 594 secondary anti-mouse (Invitrogen, Cat# A11005 1:1000), Alexa Fluor 647 conjugated streptavidin (Invitrogen, Cat# S32357 1:2500) and Dapi (1:10000) in 2% BSA for 1 hour. Cells were washed 3X with PBS and slides were mounted in ProLong Gold AntiFade (Molecular Probes, ThermoFisher Scientific, Cat# P36930) and imaged on a Nikon Eclipse Ti2 at 60X or 100X objective with oil immersion or at 20X using In Cell Analyzer 6000 microscope (GE Healthcare). Images were captured and analyzed using Elements software (v 5.41.02). For EGFR and LDLR colocalization immunofluorescence, HeLa cells stably expressing cytoplasmically tagged LDLR were serum starved overnight then stimulated with EGF for 15 minutes. Cells were then fixed and probed with EGFR antibody (Cell signaling, Cat# 4267, 1:100) and anti-FLAG antibody (1:500).

### Western and far-western blotting

After inducing expression, cells were labeled with biotin in the presence or absence of ATP and vitamin B5. Cells were then washed twice with cold PBS and lysed in modified RIPA (modRIPA: 50 mM Tris-HCl, pH 7.4, 150 mM NaCl, 1 mM EGTA, 0.5 mM EDTA, 1 mM MgCl2, 1 % NP 40, 0.1% SDS, 0.4 % sodium deoxycholate, 1 mM PMSF (Bioshop, Canada, Cat# PMS 123.5), 1x Protease Inhibitor mixture buffer (Sigma-Aldrich, Cat# P8340), 250 U of Benzonase (Sigma, Cat# 71205-3) and 1 µg of RNase A) on ice. SDS concentration was increased to 1 % after which cells were scraped and lysates were spun at 15000 g for 10 min. Proteins were then boiled in Laemmli sample buffer prepared in house and resolved on a 10 % polyacrylamide gel and transferred to nitrocellulose membranes (GE Healthcare Life Science, Uppsala, Sweden, Cat# 10600001). For bait expression detection, membranes were blocked in 3 % non-fat milk in Tris-buffered saline (TBS) with 0.1 % Tween-20 (TBST). Proteins were probed using mouse anti-FLAG antibody (Monoclonal anti-FLAG M2 antibody, Sigma-Aldrich, F3165, 1:2000), or anti-beta actin (Abcam, Cat# ab227, 1:5000), in blocking buffer, washed in TBST and detected with anti-Mouse IgG-Horseradish peroxidase (GE Healthcare Life Science, Cat# NA931, 1:5000). For streptavidin staining, nitrocellulose membranes were blocked in 3 % BSA in TBST and stained with HRP-conjugated streptavidin (GE Healthcare Life Science, Cat# RPN1231vs,1:2500) in blocking solution. Membranes were developed using LumiGLO chemiluminescent reagent (Cell Signaling Technology, Danvers, MA, Cat# 7003S) or ECL reagent (Global Life Sciences Solutions USA LLC, Cat# RPN2232) and imaged using a BioRad ChemiDoc (BioRad, Mississauga, ON, Canada).

### Biotin Labeling and Affinity Purification

BioID was performed as previously described (*60*) with some modifications. Briefly, cells were seeded in 150 mm plates and expression induced at 70 % confluence. After 24 hours, cells were labeled with 25 µM biotin in the presence or absence of ATP and vitamin B5 for 15 minutes. Cells were lysed in 1 ml of modRIPA (as explained in previous section). Lysates were collected in Eppendorf tubes, snap frozen on dry ice and stored in −80 °C until processed. Cell lysates were thawed on ice and sonicated for 15 s (5 seconds on, 3 seconds off for three cycles) at 30 % amplitude on a Q500 Sonicator with 1/8” Microtip (QSonica, Newtown, Connecticut, Cat# 4422). Samples were centrifuged at 15,000 x g for 15 min and the proteins in the supernatant were reduced and alkylated with 5 mM DTT and 20 mM IAA, respectively. Proteins (except from cells expressing eGFP and cytoplasmically tagged EGFR) were further treated with 250 U of PNGase F (NEB, Cat# P0704) for 1 hour at 37 °C. Biotinylated proteins were then captured using 15 µl (packed volume) of pre-washed Streptavidin agarose beads (GE Healthcare Life Science, Cat# 17511301). After overnight incubation at 4 °C with rotation, streptavidin beads were washed once with SDS-Wash buffer (25 mM Tris-HCl, pH 7.4, 2 % SDS), once with RIPA (50 mM Tris-HCl, pH 7.4, 150 mM NaCl, 1 mM EDTA, 1 % NP 40, 0.1 % SDS, 0.4 % sodium deoxycholate), once with TNEN buffer (25 mM Tris-HCl, pH 7.4, 150 mM NaCl, 1 mM EDTA, 0.1% NP40), and three times with 50 mM ammonium bicarbonate (ABC) buffer (pH 8.0). On-bead digestion was performed with 1 µg of trypsin (Sigma Aldrich, Cat# 6567) in 70 µl of ABC buffer, overnight with rotation at 37 °C, followed by further digestion with an additional 0.5 µg of trypsin for 3 h. Following digestion, beads were spun down (400g for 30 s), and the supernatant was transferred to a new tube. The beads were washed twice with HPLC grade water, and the washes were pooled with the peptide supernatant. A final spin was performed at 10,000 rpm for 2 min, and the supernatant was transferred to a new tube (keeping 20 µl of peptide volume at the bottom of the tube to avoid bead carryover). The peptides were acidified with 0.1 volume of 50% formic acid and dried by vacuum centrifugation. Samples were stored at −80 °C then re-suspended in 2.5 % formic acid prior to mass spectrometry analysis.

### S-trap column digestion for total protein quantification

Hela cells were cultured in 100mm plates and serum starved overnight at 80% confluence. The next day cells were stimulated with 20 ng/ml for 15 minutes followed by lysis in 5% SDS and 50 mM triethylammonium bicarbonate (TEAB) according to the manufacturer’s instructions (S-trap micro spin column digestion, ProtiFi). Briefly, 50 µg of total cell lysate was reduced, alkylated, and acidified using 2 % phosphoric acid. Proteins were then resuspended with S-trap buffer (90 % methanol, 100 mM TEAB), loaded onto columns, and washed with S-trap buffer. Proteins were in-column digested with 1:25 trypsin for 1 hour at 47 °C after which the peptides were eluted with 50 mM TEAB followed by 0.2 formic acid in 50 % acetonitrile and vacuum dried prior to analysis by MS.

### Mass Spectrometry Acquisition

Tryptic peptides from cells expressing ecTurboID-TMD (data figure 2) were analyzed on a TripleTOF 6600 (SCIEX, Framingham, MA) with a nanoelectrospray ion source connected in-line to a 425 Nano-HPLC system (Eksigent Technologies, Dublin, CA). Two thirds of each tryptic peptide sample from a 150mm cell culture plate was injected into a home packed (3 μm Reprosil-Pur 120 C18-AQ, Dr. Maisch, Ammerbuch, Germany) pulled tip fused silica capillary column (100 μm internal diameter, 20 cm length) with a 5 to 8 μm tip opening generated using a laser puller (Sutter Instrument Co, model P-2000). Samples were loaded onto the LC using an autosampler at 800 nl/min and eluted at 400 nl/min over a 90-minute gradient with solvent composition rising from 2% to 35% acetonitrile with 0.1% formic acid. The mass spectrometer was operated in a data-dependent acquisition (DDA) method using an accumulation time of 250 ms and a mass range of 400 to 1800 m/z for MS1 followed by 10 MS/MS scans each with a 100 ms accumulation time. Only ions with a charge state between 2+ and 5+ were analyzed and were then excluded for 7 seconds with a 50 mDa mass tolerance. Precursor selection used an isolation width of 0.7 m/z, and minimum intensity threshold of 200.

Tryptic peptides from cells expressing EGFR, ERBB2, CD147, LDLR, BMPR1A, eGFP, GPI and TMD tagged with ecTurboID (data figures 3 to 6) were run on a timsTOF Pro 2 quadrupole time of flight (qToF) trapped ion mobility mass spectrometer (Bruker, Billerica, Massachusetts) connected to a NanoElute liquid chromatography system (Bruker, Billerica, Massachusetts). One sixth of each tryptic peptide sample from a 150mm cell culture plate was injected onto a home packed (1.9 µm ReproSil Gold C18, Dr. Maisch, Ammerbuch, Germany) pulled tip fused silica capillary column (75 µm internal diameter, 50 cm in length). Samples were loaded onto the column at 800 bar constant pressure using the autosampler before being separated at 100 or 150 nl/min over a 90-minute gradient with solvent composition rising from 2 % to 35 % acetonitrile with 0.1 % formic acid. The mass spectrometer was operated in a DDA PASEF mode with 10 PASEF frames per cycle and active dynamic exclusion.

Precursors were selected using a polygonal filter which excluded singly charged species by their ion mobility. Ion mobility was ramped from 0.6 to 1.6 1/K0 in 100 ms, with a matching 100 ms accumulation time resulting in a total cycle time up to 1.16 s. Collision energy for the selected precursors was non-linearly dependent on ion mobility within the range of 17.12 eV at inverse mobility 0.6 to 76.46 eV at inverse mobility 1.6.

Tryptic peptides from cells expressing LDLR-N and LDLR-C (data figure 7) were also run on a timsTOF Pro 2 qToF, following the same DDA method described above, connected to Evosep One (Evosep, Odense, Denmark) liquid chromatography system using the 30 sample per day (30SPD) standard gradient and EvoSep EV1106 Performance column (15 cm x 150 μm, 1.9 μm C18 packing). The column was coupled to timsTOF via a 20 μm silica emitter (Bruker Captive Spray ZDV Sprayer 20) and held at 40 °C in a Bruker Column Toaster column oven. One sixth of each tryptic peptide sample from a 150 mm cell culture plate was loaded onto EvoTip Pure sample tips using the manufacturer recommended procedure before separation using the default 30SPD and 44-minute EvoSep method (Evosep, Odense, Denmark).

Tryptic peptides from HeLa cells digested on S-trap column were run on the timsTOF Pro 2 connected to Evosep One (Evosep, Odense, Denmark) liquid chromatography system using the 60 sample per day (60SPD) standard gradient and EvoSep EV1109 Performance column (8 cm x 150 μm, 1.5 μm C18 packing). 250 ng of tryptic peptides were loaded onto EvoTip Pure sample tips using the manufacturer recommended procedure before separation using the default 60SPD and 22-minute EvoSep method (Evosep, Odense, Denmark). Peptides were run in both PASEF and data independent acquisition (DIA), diaPASEF modes. DIA was performed on a separate injection of the sample with the same separation conditions as DDA but with diaPASEF MS acquisition. The diaPASEF was set up with 20 windows per cycle each covering a 50 m/z slice running from 400 m/z to 1350 m/z with a 1 m/z overlap between windows. The windows cover the entirety of the 0.85 to 1.3 inverse mobility range. The total cycle time was approximately 1.05 seconds. DIA collision energy was selected based on the precursor mobility ranging from 29.56 eV at inverse mobility 0.85 to 58.09 eV at inverse mobility 1.3.

### Mass Spectrometry Data Analysis

Mass spectrometry data files were stored and searched using ProHits laboratory information management system (LIMS) platform (*61*). ProteoWizard (V3.0.1072) (*62*) was used to convert the .RAW files from 6600 TOF to. mgf and. mzML formats. These data files were searched using Mascot (V2.3.02) (*63*) and Comet (V2016.01 rev.2) (*64*) against human RefSeq database (version 57, September 12,2020) with bovine sequences from the RefSeq database (version 201, September 12, 2020), supplemented with “common contaminants” from the Max Planck Institute (http://www.coxdocs. org/doku.php?id=maxquant: start_downloads.htm) and the Global Proteome Machine (GPM; ftp://ftp.thegpm.org/fasta/cRAP/crap.fasta) with sequence tags (TurboID, BirA, GST26, mCherry and GFP), LysC, and streptavidin. The total number of entries including reverse (decoy) sequences was 200288. The search was set to identify tryptic peptides allowing for 2 missed cleavages per peptide and a mass tolerance of 35 ppm for 2+ to 4+ peptides. Fragment ion mass tolerance was set to 0.15 Da. Carbamidomethylation (C) was set as a fixed modification while deamidation and oxidation (Met) were selected as variable modifications.

Data files from timsTOF were searched using Fragpipe (version 17) within ProHits, directly from the Bruker ‘.d’ format against human Uniprot database (UP000005640, October 26, 2021) supplemented with “common contaminants” from the Global Proteome Machine (GPM; ftp://ftp.thegpm.org/fasta/cRAP/crap.fasta) with sequence tags (BirA, TurboID, miniTurbo, GST26, mCherry, GFP, eGFP) excluding the human proteins. The total number of entries was 40841 including decoys. The search was set to identify tryptic peptides allowing for 2 missed cleavages per peptide.

Peptide mass tolerance was set to ± 20 ppm and fragment ion mass tolerance was ± 20 ppm. Peptide fixed and variable modifications were set as mentioned above in addition to Acetyl (Protein N-term) as a variable modification. Results from each search engine were analyzed through the Trans-Proteomic Pipeline (TPP, v.4.7 POLAR VORTEX rev 1) (*65*) using iProphet (*66*). Results from Mascot and Comet were combined prior to analysis by TPP. All proteins with an iProphet probability ≥ 95% were used for analysis. For analyzing diaPASEF data files, a spectral library was built from DDA files with the same search parameters listed above using Fragpipe within Prohits. The spectral library was then used to search DIA samples using DiaNN 1.8.2 (*67*). Default search parameters were used including match between runs except the quantitation strategy was “Robust LC” and protein inference was disabled. All identified proteins were validated using 1% FDR for both DDA and DIA data.

### SAINT analysis

SAINTexpress (Version 3.6.1)(*47*) was used to score the probability of high confidence proximity interactions. SAINTexpress is based on scoring the interaction of an identified prey with a specific bait in comparison to controls based on spectral counting for each prey. Each bait, condition and construct were profiled using purifications from two independent labelling experiments. For LDLR follow-up experiments with and without EGF stimulation, triplicate purifications were analyzed. Controls were composed of prey lists obtained from HeLa cells expressing eGFP tagged with TurboID (and cultured in complete growth media) and prey lists of baits profiled in the absence of ATP. The latter generates a list of proteins that interact with the bait intracellularly, not at the cell membrane. Each SAINTexpress analysis included 4 eGFP runs and all “no ATP” runs from the corresponding baits being compared, comprising a minimum of 8 controls within each dataset. For LDLR proximity interaction profiling with and without EGF, HeLa cells expressing TurboID-tagged eGFP were serum starved overnight and prey lists were used as controls for more stringency in scoring interactions. Scores were averaged across biological replicates, and these averages were used to calculate a Bayesian False Discovery Rate (BFDR); preys detected with a BFDR of ≤ 1% were considered high confidence interactors.

### Data Visualization

Bait-prey dot plots, heatmaps, and bait versus bait (or condition versus condition) plots were generated using ProHits-viz (http://profits-viz.org (*68*)). The volcano plot was generated in R from text files downloaded from DiaNN searches with protein intensity values, excluding proteins with missing values in at least 1 replicate. GO enrichment analysis was done in g:Profiler (*69*) either within ProHits-viz or directly from the website using default parameters (user threshold was set to 0.01) and the human database. Data was downloaded as csv files, with the Padj (g:Profiler default) and number of genes in a GO term included. Enrichment dot plots were generated using in-house developed codes in Python (available on Github at https://github.com/gingraslab/go_terms_dot_plot) that visualize the top enriched Gene Ontology (GO) terms (the top 5 terms for each category assessed were plotted here) of a group of genes from different conditions, producing a dot plot that facilitates analysis and comparison across conditions. Bar graphs were generated using GraphPad (v 9.5.1) and Microsoft Excel. Venn diagrams were created using DeepVenn (www.deepvenn.com). Cartoon diagrams were created with BioRender (www.biorender.com). All images were adapted for publication using adobe illustrator (v 27.7). Fluorescent images taken on the InCell analyzer 6000 microscope were captured at 20X, with 16 images taken per condition then stitched to make 1 image using an in-house developed code in Matlab. Intensity quantification was done in Columbus.

In ProHits-viz dot plot tool, once a prey passes the selected BFDR threshold (≤1% used here) with one bait, all its spectral count values across all baits are shown. The BFDR of the prey is then indicated by the edge color while spectral count is represented by the color gradient of the node. The size of the node indicates the relative prey counts across all baits in reference to the bait in which it was detected with the highest number of spectral counts. In the GO enrichment dot plot tool, the color of the circle refers to the p-value of enrichment generated by g:Profiler and the size of the circle refers to the number of genes in our dataset contributing to the enriched term.

## Acknowledgments

We thank Julia Kitaygorodsky for generating panel A of supplementary figure 7, U. Dionne for comments and valuable discussions, Tess Branon and Alice Ting for the TurboID sequence information and members of the Gingras lab for sharing reagents and information. We further thank members of the proteomics core of the Network Biology Collaborative Centre and the OPTical IMAging (OPTIMA) Facility at the Lunenfeld-Tanenbaum Research Institute for technical advice and support.

## Author contributions

R.A.M., P.S.T and A.C.G. conceived the study. P.S.T. designed constructs and performed initial experiments. R.A.M. performed all experiments presented in the figures. B.S. developed the mass spectrometry analysis pipeline and helped analyze the samples. V.K. developed visualization tools. C.E.M. provided technical support and participated in data analysis and discussion writing. R.A.M. and A.C.G. wrote the manuscript with support by U. Dionne, and all authors approved the final manuscript.

## Data and materials availability

Proteomics data has been submitted to ProteomeXchange via partner MassIVE (massive.ucsd.edu) with the following identifiers PXD046334 and MSV000093178 and available at ftp://MSV000093178@massive.ucsd.edu.

---

**Figure S1:**
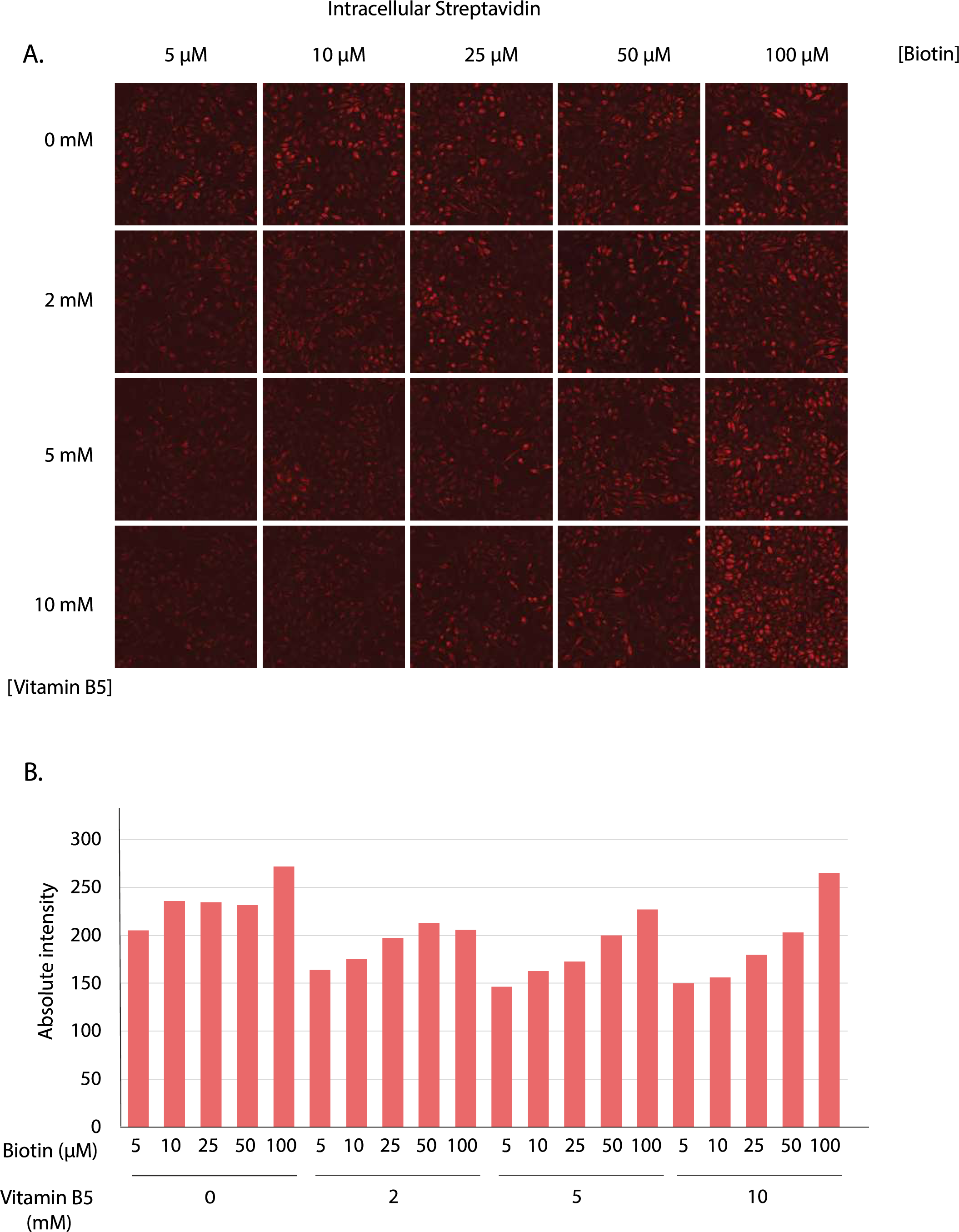
Establishing ecTurboID labeling concentrations 1. (**A**) HeLa cells stably expressing ecTurboID-TMD and treated for 30 minutes with the indicated concentrations of biotin and vitamin B5 in the presence of 1.5 mM ATP were fixed and stained to show intracellular labeling as illustrated in the bottom left cartoon in panel A of figure 1. Images are representative of 16 fields taken for each condition at 20X and stitched together into one image. (**B**) Intensity quantification of intracellular labeling from images in panel A.

**Figure S2:**
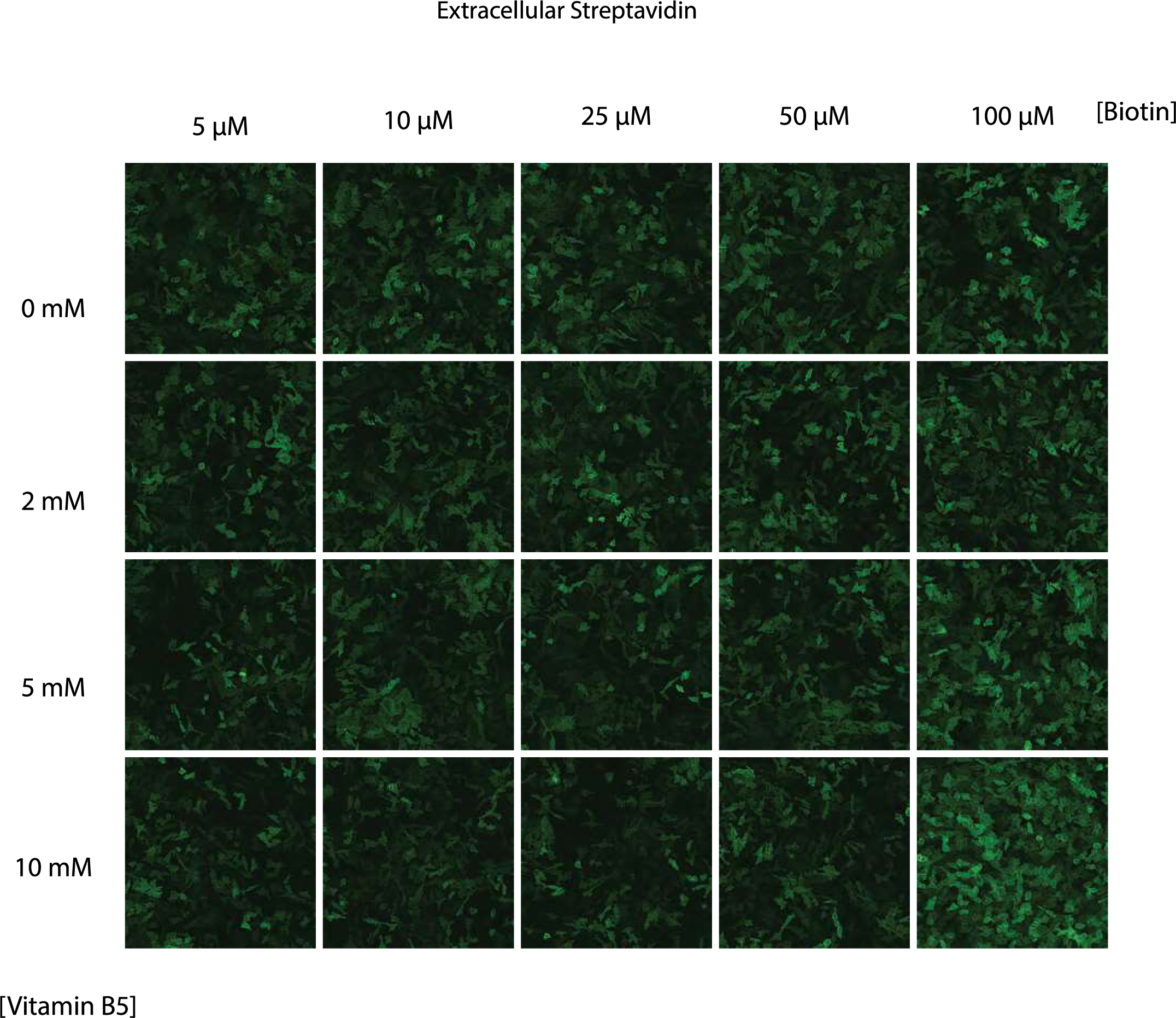
Establishing ecTurboID labeling concentrations 2. HeLa cells stably expressing ecTurboID-TMD and treated for 30 minutes with the indicated concentrations of biotin and vitamin B5 in the presence of 1.5 mM ATP were fixed and stained to show extracellular labeling as illustrated in the bottom left cartoon in panel A of Figure 1.

**Figure S3:**
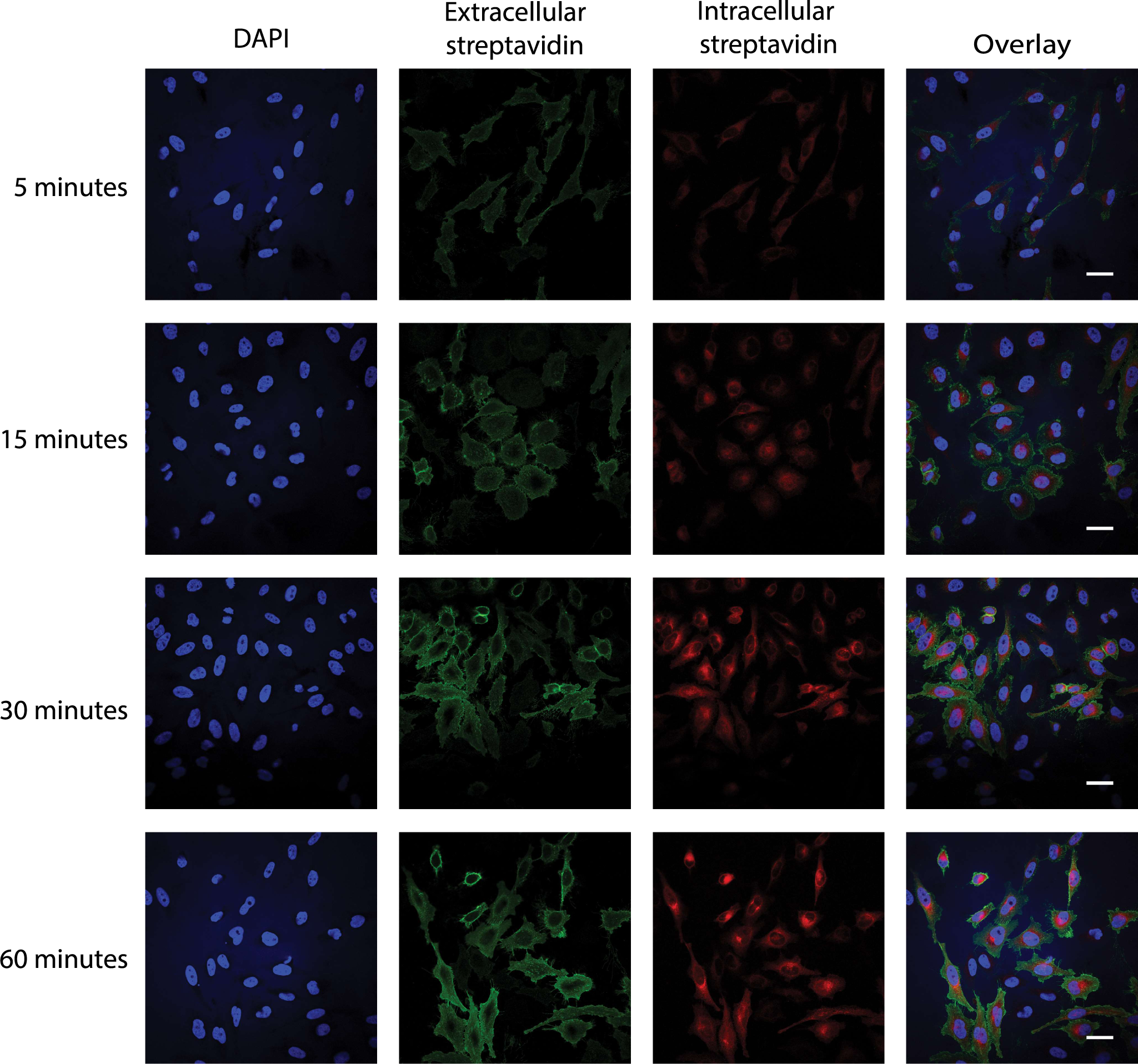
Establishing ecTurboID labeling time. HeLa cells stably expressing ecTurboID-TMD were labeled with 25 µM biotin for 5, 15, 30 and 60 minutes in the presence of 5 mM vitamin B5 and 1.5 mM ATP. Cells were stained to show extracellular labeling (green fluorescence) and intracellular labeling (red fluorescence), as illustrated in the bottom left cartoon in panel A of Figure 1. Scale bar, 20 µm.

**Figure S4:**
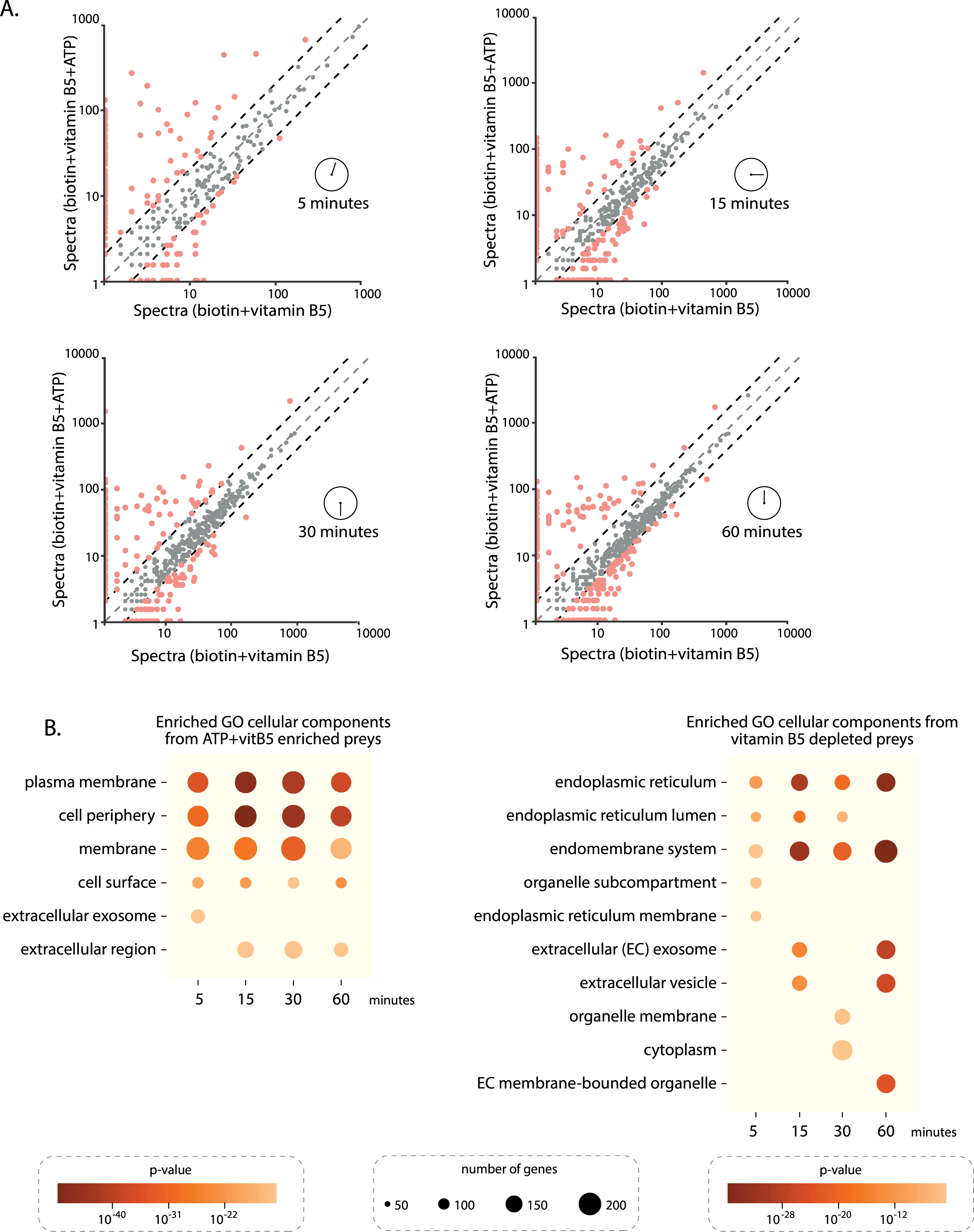
Profiling ecTurboID-TMD at different labeling time points. (**A**) Condition versus condition scatter plots of abundance (average spectral counts) of all proteins identified at 5, 15, 30 and 60 minutes with 25 µM biotin and 5 mM vitamin B5 in the presence or absence of 1.5 mM ATP in HeLa cells expressing ecTurboID-TMD. (**B**) Gene Ontology (GO) analysis of cellular components enriched from all proteins with a 2-fold change (enriched or depleted) or more identified at 5, 15, 30 and 60 minutes with biotin and vitamin B5 (vitB5) in the absence or presence of ATP (same concentrations listed in A).

**Figure S5:**
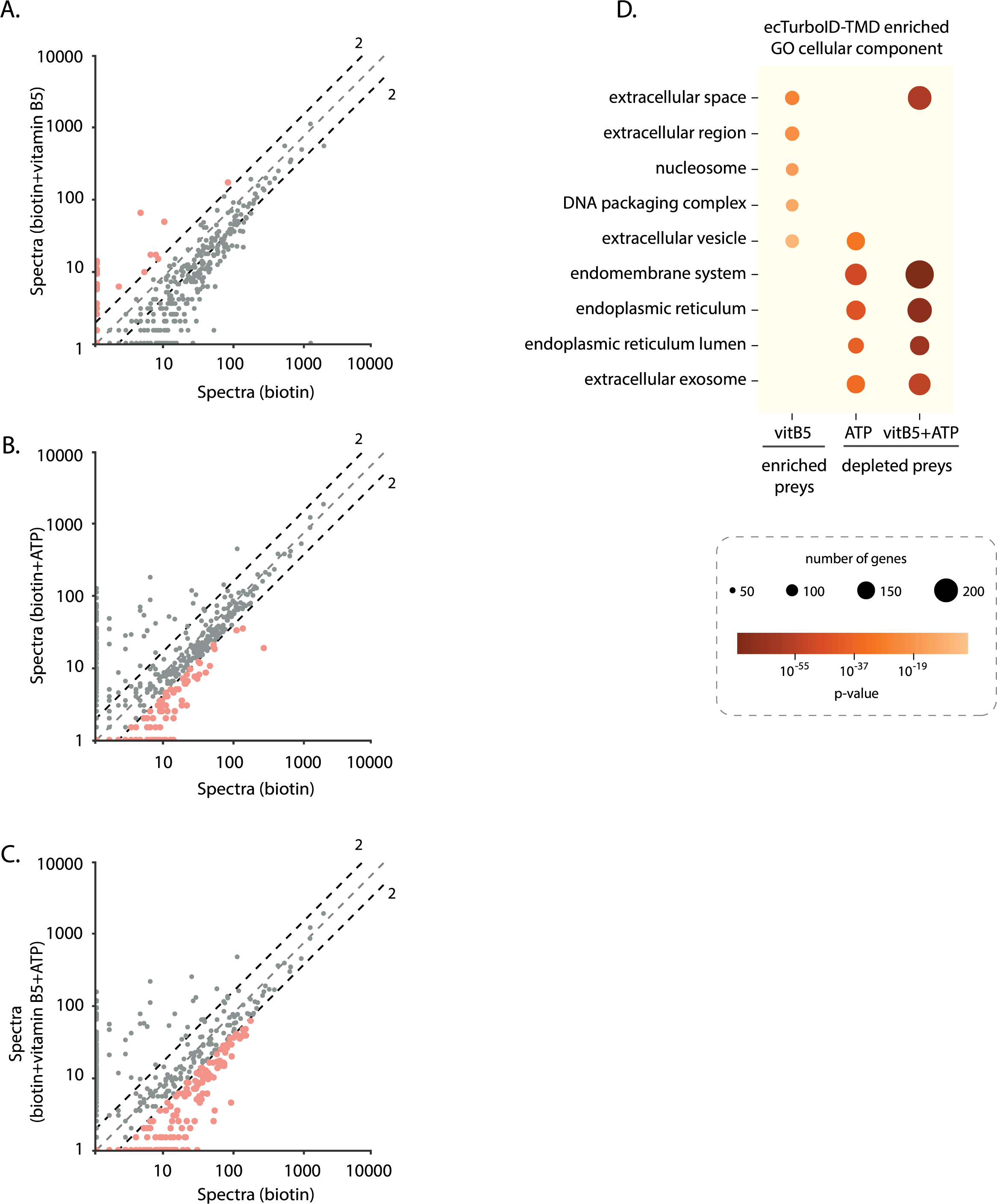
Gene Ontology analysis of ecTurboID-TMD preys enriched with vitamin B5 and depleted with ATP. (**A–C**) Condition versus condition scatter plots of abundance (average spectral counts) of all proximity interactors identified with 15 minutes of 25 µM biotin labeling in the presence of 5 mM vitamin B5 or 1.5 mM ATP or both in HeLa cells expressing ecTurboID-TMD (same data set analyzed in Figure 2) (**D**) Gene Ontology (GO) analysis of cellular components enriched and depleted from proximity interactors with a 2-fold change or more upon addition of vitamin B5 (vitB5), ATP or both.

**Figure S6:**
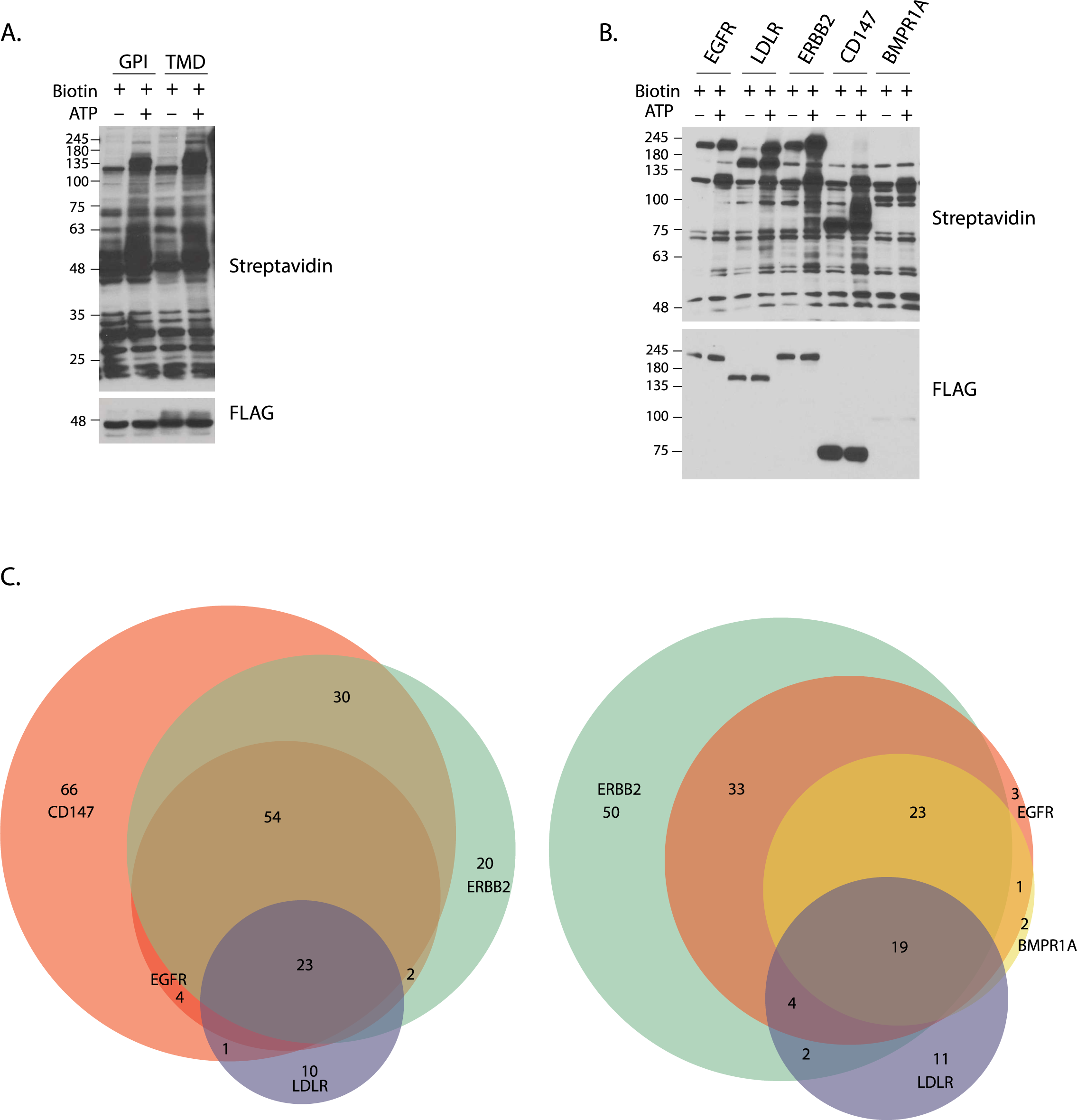
Comparing constructs and tagged proteins. (**A**) Lysates from HeLa cells expressing ecTurboID-TMD, and ecTurboID-GPI labeled for 15 minutes with 25 µM biotin in the presence or absence of 1.5 mM ATP were probed for expression (anti-FLAG) and biotinylation (HRP-conjugated streptavidin). (**B**) Lysates from HeLa cells expressing EGFR, LDLR, ERBB2, CD147, and BMPR1A fused in frame with ecTurboID labeled for 15 minutes with 25 µM biotin in the presence or absence of 1.5mM ATP were probed for expression (anti-FLAG) and biotinylation (HRP-conjugated streptavidin). (**C**) Venn diagram of all high confidence proximity interactors (BFDR≤ 1% and minimum of 2 spectral counts) identified in HeLa cells expressing EGFR, ERBB2, CD147, LDLR and BMPR1A fused in frame with the ecTurboID labeled for 15 minutes in the presence of 1.5 mM ATP and 5 mM vitamin B5.

**Figure S7:**
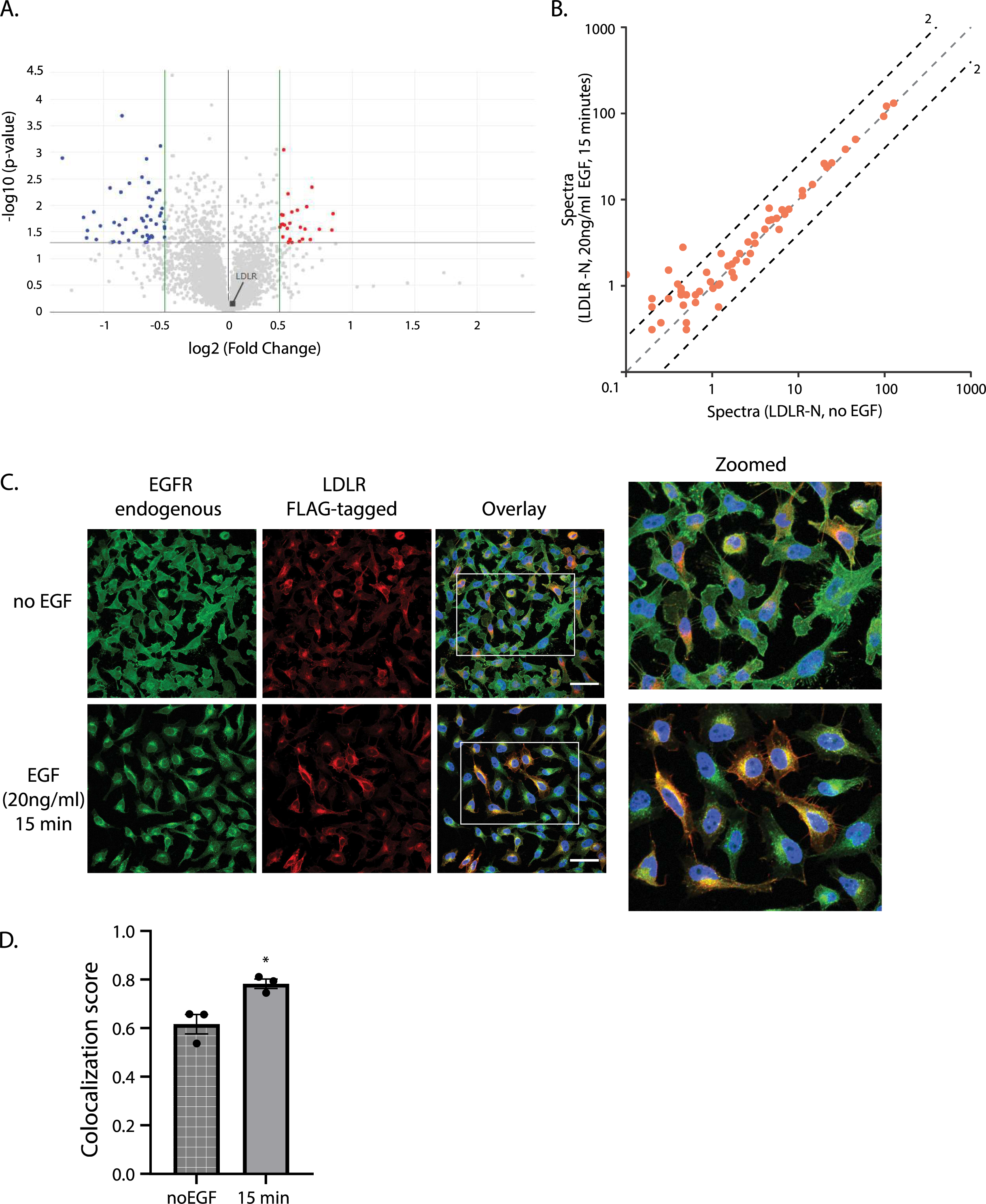
Characterization of LDLR-EGFR association. (**A**) Volcano plot showing total proteome change in HeLa cells after 15 minutes of 20 ng/ml EGF stimulation compared to resting state. The x-axis shows log2 fold change, based on intensity quantification, marked by green lines at 2-fold cutoff and the y-axis shows -log10 p-value from unpaired t-test marked by gray line at 0.05 cut off, *n* = 3. (**B**) Condition versus condition scatter plot of abundance (average spectral counts plotted as log10 values) for all high confidence proximity interactors (BFDR≤ 0.01 and minimum of 2 spectral counts) identified after 15 minutes of 25 µM biotin, 5 mM vitamin B5 and 1.5mM ATP in LDLR-N expressing HeLa cells in the presence or absence of 20 ng/ml EGF. (**C**) HeLa cells stably expressing LDLR-C were stimulated with 20 ng/ml EGF for 15 minutes, then fixed and probed for endogenous EGFR and exogenous LDLR using anti-FLAG antibody. (**D**) Pearson colocalization score from a minimum of ninety cells in three fields randomly selected for each condition. Values are mean ± standard error of the mean for Pearson coefficient generated from NIS Elements software (v 5.30.01) colocalization analysis (*P <0.05, two-tailed paired *t-test*).

**Table S1:**
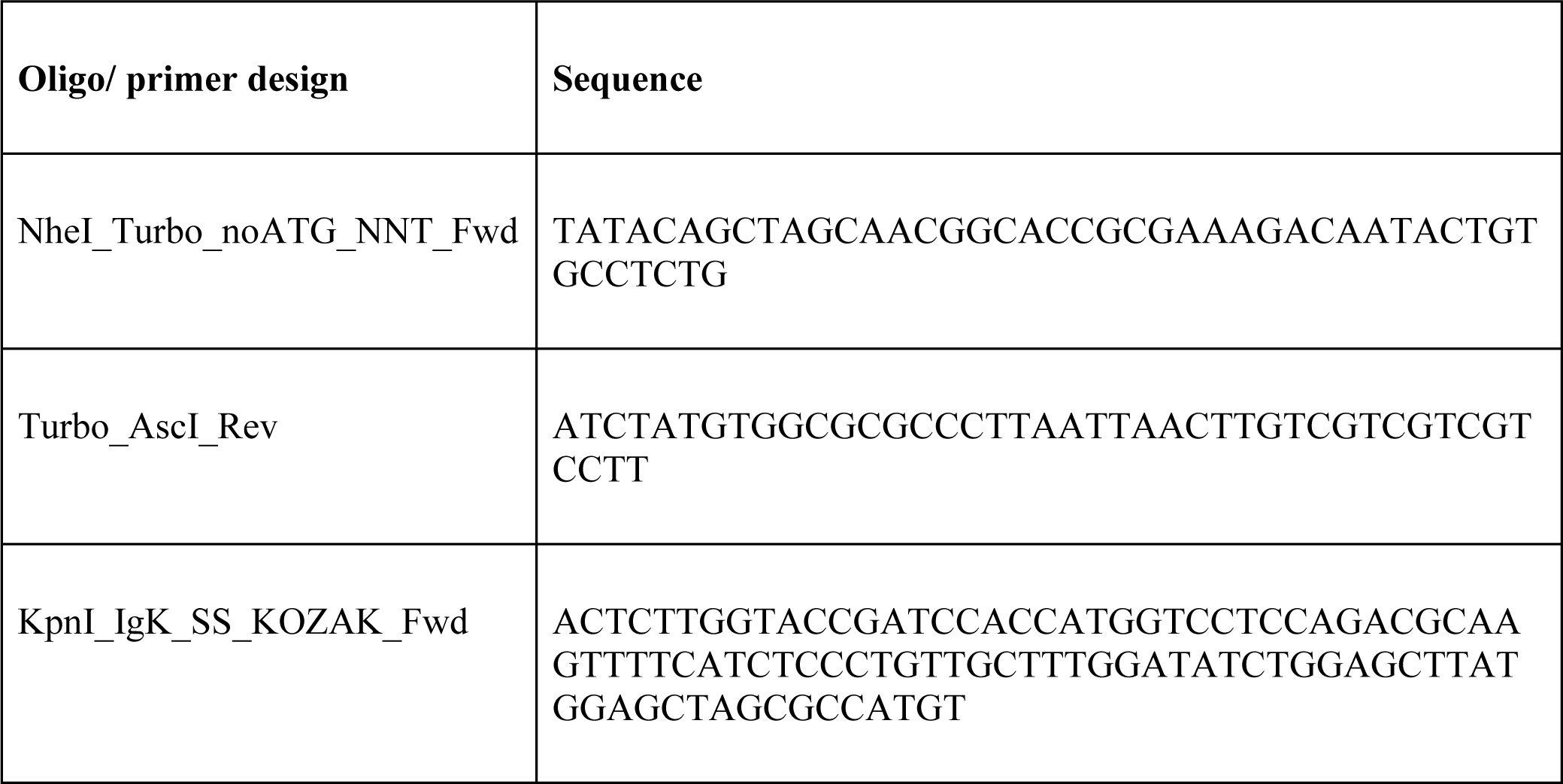
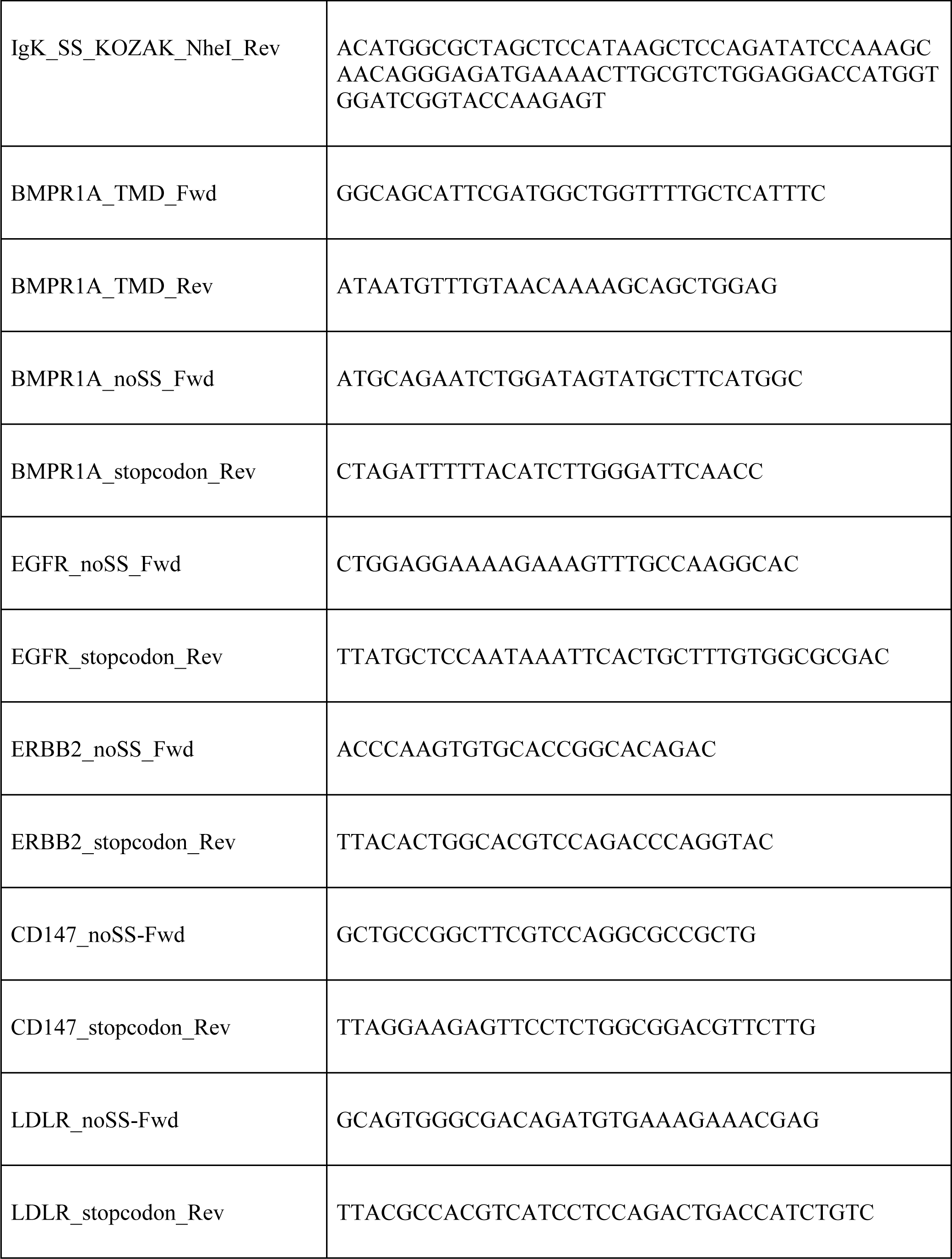
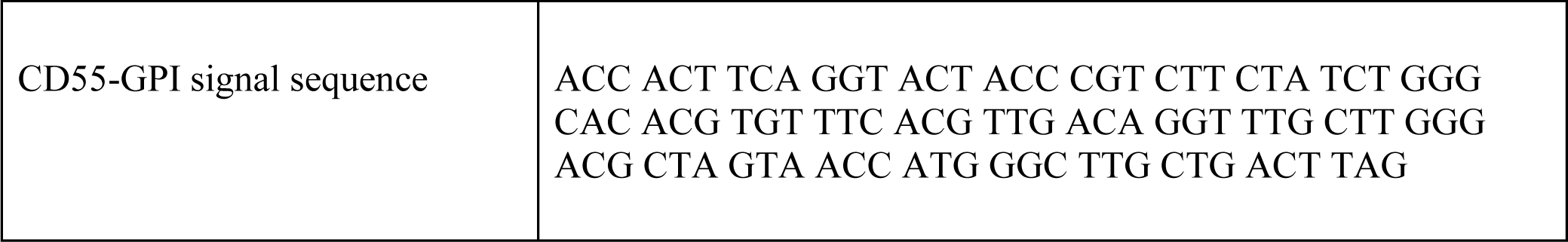
Oligo and primer sequences.

## REFERENCES

1. D. Bausch-Fluck, E. S. Milani, B. Wollscheid, Surfaceome nanoscale organization and extracellular interaction networks. Curr. Opin. Chem. Biol. 48, 26–33 (2019).

2. K. Jacobson, P. Liu, B. C. Lagerholm, The Lateral Organization and Mobility of Plasma Membrane Components. Cell. 177, 806–819 (2019).

3. M. R. Chastney, J. R. W. Conway, J. Ivaska, Integrin adhesion complexes. Curr. Biol. 31, R536– R542 (2021).

4. E. S. Novoseletskaya, P. V. Evdokimov, A. Y. Efimenko, Extracellular matrix-induced signaling pathways in mesenchymal stem/stromal cells. Cell Commun. Signal. 21, 244 (2023).

5. A. Tomas, C. E. Futter, E. R. Eden, EGF receptor trafficking: consequences for signaling and cancer. Trends Cell Biol. 24, 26–34 (2014).

6. Z. J. Gartner, J. A. Prescher, L. D. Lavis, Unraveling cell-to-cell signaling networks with chemical biology. Nat. Chem. Biol. 13, 564–568 (2017).

7. A.-C. Gingras, K. T. Abe, B. Raught, Getting to know the neighborhood: using proximity-dependent biotinylation to characterize protein complexes and map organelles. Curr. Opin. Chem. Biol. 48, 44–54 (2019).

8. M. M. Makowski, E. Willems, P. W. T. C. Jansen, M. Vermeulen, Cross-linking immunoprecipitation-MS (xIP-MS): Topological Analysis of Chromatin-associated Protein Complexes Using Single Affinity Purification. Mol. Cell. Proteomics. 15, 854–865 (2016).

9. J. Zheng, Z. Zheng, C. Fu, Y. Weng, A. He, X. Ye, W. Gao, R. Tian, Deciphering intercellular signaling complexes by interaction-guided chemical proteomics. Nat. Commun. 14, 4138 (2023).

10. Z. Hakhverdyan, M. Domanski, L. E. Hough, A. A. Oroskar, A. R. Oroskar, S. Keegan, D. J. Dilworth, K. R. Molloy, V. Sherman, J. D. Aitchison, D. Fenyö, B. T. Chait, T. H. Jensen, M. P. Rout, J. LaCava, Rapid, optimized interactomic screening. Nat. Methods. 12, 553–560 (2015).

11. J. LaCava, K. R. Molloy, M. S. Taylor, M. Domanski, B. T. Chait, M. P. Rout, Affinity proteomics to study endogenous protein complexes: pointers, pitfalls, preferences and perspectives. Biotechniques. 58, 103–119 (2015).

12. S. Heo, S. Spoerk, R. Birner-Gruenberger, G. Lubec, Gel-based mass spectrometric analysis of hippocampal transmembrane proteins using high resolution LTQ Orbitrap Velos Pro. Proteomics. 14, 2084–2088 (2014).

13. C. S. Müller, W. Bildl, A. Haupt, L. Ellenrieder, T. Becker, C. Hunte, B. Fakler, U. Schulte, Cryo-slicing Blue Native-Mass Spectrometry (csBN-MS), a Novel Technology for High Resolution Complexome Profiling*. Mol. Cell. Proteomics. 15, 669–681 (2016).

14. J. A. Bosch, C.-L. Chen, N. Perrimon, Proximity-dependent labeling methods for proteomic profiling in living cells: An update. Wiley Interdiscip. Rev. Dev. Biol. 10, e392 (2021).

15. N. St-Denis, G. D. Gupta, Z. Y. Lin, B. Gonzalez-Badillo, A. O. Veri, J. D. R. Knight, D. Rajendran, A. L. Couzens, K. W. Currie, J. M. Tkach, S. W. T. Cheung, L. Pelletier, A.-C. Gingras, Phenotypic and Interaction Profiling of the Human Phosphatases Identifies Diverse Mitotic Regulators. Cell Rep. 17, 2488–2501 (2016).

16. M. Akdag, Z. S. Yunt, A. Kamacioglu, M. H. Qureshi, B. A. Akarlar, N. Ozlu, Proximal Biotinylation-Based Combinatory Approach for Isolating Integral Plasma Membrane Proteins. J. Proteome Res. 19, 3583–3592 (2020).

17. B. T. Lobingier, R. Hüttenhain, K. Eichel, K. B. Miller, A. Y. Ting, M. von Zastrow, N. J. Krogan, An Approach to Spatiotemporally Resolve Protein Interaction Networks in Living Cells. Cell. 169, 350–360.e12 (2017).

18. C. D. Go, J. D. R. Knight, A. Rajasekharan, B. Rathod, G. G. Hesketh, K. T. Abe, J.-Y. Youn, P. Samavarchi-Tehrani, H. Zhang, L. Y. Zhu, E. Popiel, J.-P. Lambert, É. Coyaud, S. W. T. Cheung, D. Rajendran, C. J. Wong, H. Antonicka, L. Pelletier, A. F. Palazzo, E. A. Shoubridge, B. Raught, A.-C. Gingras, A proximity-dependent biotinylation map of a human cell. Nature. 595, 120–124 (2021).

19. K. J. Roux, D. I. Kim, M. Raida, B. Burke, A promiscuous biotin ligase fusion protein identifies proximal and interacting proteins in mammalian cells. J. Cell Biol. 196, 801–810 (2012).

20. T. C. Branon, J. A. Bosch, A. D. Sanchez, N. D. Udeshi, T. Svinkina, S. A. Carr, J. L. Feldman, N. Perrimon, A. Y. Ting, Efficient proximity labeling in living cells and organisms with TurboID. Nat. Biotechnol. 36, 880–887 (2018).

21. D. I. Kim, K. C. Birendra, W. Zhu, K. Motamedchaboki, V. Doye, K. J. Roux, Probing nuclear pore complex architecture with proximity-dependent biotinylation. Proc. Natl. Acad. Sci. U. S. A. 111, E2453–61 (2014).

22. A. L. Couzens, J. D. R. Knight, M. J. Kean, G. Teo, A. Weiss, W. H. Dunham, Z.-Y. Lin, R. D. Bagshaw, F. Sicheri, T. Pawson, J. L. Wrana, H. Choi, A.-C. Gingras, Protein interaction network of the mammalian Hippo pathway reveals mechanisms of kinase-phosphatase interactions. Sci. Signal. 6, rs15 (2013).

23. G. G. Hesketh, F. Papazotos, J. Pawling, D. Rajendran, J. D. R. Knight, S. Martinez, M. Taipale, D. Schramek, J. W. Dennis, A.-C. Gingras, The GATOR–Rag GTPase pathway inhibits mTORC1 activation by lysosome-derived amino acids. Science. 370, 351–356 (2020).

24. J.-Y. Youn, W. H. Dunham, S. J. Hong, J. D. R. Knight, M. Bashkurov, G. I. Chen, H. Bagci, B. Rathod, G. MacLeod, S. W. M. Eng, S. Angers, Q. Morris, M. Fabian, J.-F. Côté, A.-C. Gingras, High-Density Proximity Mapping Reveals the Subcellular Organization of mRNA-Associated Granules and Bodies. Mol. Cell. 69, 517–532.e11 (2018).

25. G. D. Gupta, É. Coyaud, J. Gonçalves, B. A. Mojarad, Y. Liu, Q. Wu, L. Gheiratmand, D. Comartin, J. M. Tkach, S. W. T. Cheung, M. Bashkurov, M. Hasegan, J. D. Knight, Z.-Y. Lin, M. Schueler, F. Hildebrandt, J. Moffat, A.-C. Gingras, B. Raught, L. Pelletier, A Dynamic Protein Interaction Landscape of the Human Centrosome-Cilium Interface. Cell. 163, 1484–1499 (2015).

26. L.-T. Wang, M.-È. Proulx, A. D. Kim, V. Lelarge, L. McCaffrey, A proximity proteomics screen in three-dimensional spheroid cultures identifies novel regulators of lumen formation. Sci. Rep. 11, 22807 (2021).

27. Z. Guo, L. J. Neilson, H. Zhong, P. S. Murray, S. Zanivan, R. Zaidel-Bar, E-cadherin interactome complexity and robustness resolved by quantitative proteomics. Sci. Signal. 7, rs7 (2014).

28. N. Kotani, J. Gu, T. Isaji, K. Udaka, N. Taniguchi, K. Honke, Biochemical visualization of cell surface molecular clustering in living cells. Proc. Natl. Acad. Sci. U. S. A. 105, 7405–7409 (2008).

29. X.-W. Li, J. S. Rees, P. Xue, H. Zhang, S. W. Hamaia, B. Sanderson, P. E. Funk, R. W. Farndale, K. S. Lilley, S. Perrett, A. P. Jackson, New insights into the DT40 B cell receptor cluster using a proteomic proximity labeling assay. J. Biol. Chem. 289, 14434–14447 (2014).

30. D. Z. Bar, K. Atkatsh, U. Tavarez, M. R. Erdos, Y. Gruenbaum, F. S. Collins, Biotinylation by antibody recognition-a method for proximity labeling. Nat. Methods. 15, 127–133 (2018).

31. S. Jiang, N. Kotani, T. Ohnishi, A. Miyagawa-Yamguchi, M. Tsuda, R. Yamashita, Y. Ishiura, K. Honke, A proteomics approach to the cell-surface interactome using the enzyme-mediated activation of radical sources reaction. Proteomics. 12, 54–62 (2012).

32. A. Miyagawa-Yamaguchi, N. Kotani, K. Honke, Expressed glycosylphosphatidylinositol-anchored horseradish peroxidase identifies co-clustering molecules in individual lipid raft domains. PLoS One. 9, e93054 (2014).

33. N. Kotani, T. Nakano, Y. Ida, R. Ito, M. Hashizume, A. Yamaguchi, M. Seo, T. Araki, Y. Hojo, K. Honke, T. Murakoshi, Analysis of lipid raft molecules in the living brain slices. Neurochem. Int. 119, 140–150 (2018).

34. K. H. Loh, P. S. Stawski, A. S. Draycott, N. D. Udeshi, E. K. Lehrman, D. K. Wilton, T. Svinkina, T. J. Deerinck, M. H. Ellisman, B. Stevens, S. A. Carr, A. Y. Ting, Proteomic Analysis of Unbounded Cellular Compartments: Synaptic Clefts. Cell. 166, 1295–1307.e21 (2016).

35. J. Li, S. Han, H. Li, N. D. Udeshi, T. Svinkina, D. R. Mani, C. Xu, R. Guajardo, Q. Xie, T. Li, D. J. Luginbuhl, B. Wu, C. N. McLaughlin, A. Xie, P. Kaewsapsak, S. R. Quake, S. A. Carr, A. Y. Ting, L. Luo, Cell-Surface Proteomic Profiling in the Fly Brain Uncovers Wiring Regulators. Cell. 180, 373–386.e15 (2020).

36. Y. Li, Y. Wang, Y. Yao, J. Lyu, Q. Qiao, J. Mao, Z. Xu, M. Ye, Rapid Enzyme-Mediated Biotinylation for Cell Surface Proteome Profiling. Anal. Chem. 93, 4542–4551 (2021).

37. L. L. Kirkemo, S. K. Elledge, J. Yang, J. R. Byrnes, J. E. Glasgow, R. Blelloch, J. A. Wells, Cell-surface tethered promiscuous biotinylators enable comparative small-scale surface proteomic analysis of human extracellular vesicles and cells. Elife. 11 (2022), doi:10.7554/eLife.73982.

38. S. S. Lam, J. D. Martell, K. J. Kamer, T. J. Deerinck, M. H. Ellisman, V. K. Mootha, A. Y. Ting, Directed evolution of APEX2 for electron microscopy and proximity labeling. Nat. Methods. 12, 51–54 (2015).

39. Q. Liu, J. Zheng, W. Sun, Y. Huo, L. Zhang, P. Hao, H. Wang, M. Zhuang, A proximity-tagging system to identify membrane protein-protein interactions. Nat. Methods. 15, 715–722 (2018).

40. O. Shafraz, B. Xie, S. Yamada, S. Sivasankar, Mapping transmembrane binding partners for E-cadherin ectodomains. Proc. Natl. Acad. Sci. U. S. A. 117, 31157–31165 (2020).

41. T. Takano, J. T. Wallace, K. T. Baldwin, A. M. Purkey, A. Uezu, J. L. Courtland, E. J. Soderblom, T. Shimogori, P. F. Maness, C. Eroglu, S. H. Soderling, Chemico-genetic discovery of astrocytic control of inhibition in vivo. Nature. 588, 296–302 (2020).

42. D. I. Kim, S. C. Jensen, K. A. Noble, B. Kc, K. H. Roux, K. Motamedchaboki, K. J. Roux, An improved smaller biotin ligase for BioID proximity labeling. Mol. Biol. Cell. 27, 1188–1196 (2016).

43. S.-S. Gao, R. Shi, J. Sun, Y. Tang, Z. Zheng, J.-F. Li, H. Li, J. Zhang, Q. Leng, J. Xu, X. Chen, J. Zhao, M.-S. Sy, L. Feng, C. Li, GPI-anchored ligand-BioID2-tagging system identifies Galectin-1 mediating Zika virus entry. iScience. 25, 105481 (2022).

44. J. V. Greiner, T. Glonek, Intracellular ATP Concentration and Implication for Cellular Evolution. Biology. 10 (2021), doi:10.3390/biology10111166.

45. Y. Uchida, K. Ito, S. Ohtsuki, Y. Kubo, T. Suzuki, T. Terasaki, Major involvement of Na(+) - dependent multivitamin transporter (SLC5A6/SMVT) in uptake of biotin and pantothenic acid by human brain capillary endothelial cells. J. Neurochem. 134, 97–112 (2015).

46. H. Choi, B. Larsen, Z.-Y. Lin, A. Breitkreutz, D. Mellacheruvu, D. Fermin, Z. S. Qin, M. Tyers, A.-C. Gingras, A. I. Nesvizhskii, SAINT: probabilistic scoring of affinity purification–mass spectrometry data. Nat. Methods. 8, 70–73 (2010).

47. G. Teo, G. Liu, J. Zhang, A. I. Nesvizhskii, A.-C. Gingras, H. Choi, SAINTexpress: improvements and additional features in Significance Analysis of INTeractome software. J. Proteomics. 100, 37– 43 (2014).

48. M. Uhlén, E. Björling, C. Agaton, C. A.-K. Szigyarto, B. Amini, E. Andersen, A.-C. Andersson, P. Angelidou, A. Asplund, C. Asplund, L. Berglund, K. Bergström, H. Brumer, D. Cerjan, M. Ekström, A. Elobeid, C. Eriksson, L. Fagerberg, R. Falk, J. Fall, M. Forsberg, M. G. Björklund, K. Gumbel, A. Halimi, I. Hallin, C. Hamsten, M. Hansson, M. Hedhammar, G. Hercules, C. Kampf, K. Larsson, M. Lindskog, W. Lodewyckx, J. Lund, J. Lundeberg, K. Magnusson, E. Malm, P. Nilsson, J. Odling, P. Oksvold, I. Olsson, E. Oster, J. Ottosson, L. Paavilainen, A. Persson, R. Rimini, J. Rockberg, M. Runeson, A. Sivertsson, A. Sköllermo, J. Steen, M. Stenvall, F. Sterky, S. Strömberg, M. Sundberg, H. Tegel, S. Tourle, E. Wahlund, A. Waldén, J. Wan, H. Wernérus, J. Westberg, K. Wester, U. Wrethagen, L. L. Xu, S. Hober, F. Pontén, A human protein atlas for normal and cancer tissues based on antibody proteomics. Mol. Cell. Proteomics. 4, 1920–1932 (2005).

49. N. Bag, S. Huang, T. Wohland, Plasma Membrane Organization of Epidermal Growth Factor Receptor in Resting and Ligand-Bound States. Biophys. J. 109, 1925–1936 (2015).

50. R. Oughtred, J. Rust, C. Chang, B.-J. Breitkreutz, C. Stark, A. Willems, L. Boucher, G. Leung, N. Kolas, F. Zhang, S. Dolma, J. Coulombe-Huntington, A. Chatr-Aryamontri, K. Dolinski, M. Tyers, The BioGRID database: A comprehensive biomedical resource of curated protein, genetic, and chemical interactions. Protein Sci. 30, 187–200 (2021).

51. D. Guo, F. Reinitz, M. Youssef, C. Hong, D. Nathanson, D. Akhavan, D. Kuga, A. N. Amzajerdi, H. Soto, S. Zhu, I. Babic, K. Tanaka, J. Dang, A. Iwanami, B. Gini, J. Dejesus, D. D. Lisiero, T. T. Huang, R. M. Prins, P. Y. Wen, H. I. Robins, M. D. Prados, L. M. Deangelis, I. K. Mellinghoff, M. P. Mehta, C. D. James, A. Chakravarti, T. F. Cloughesy, P. Tontonoz, P. S. Mischel, An LXR agonist promotes glioblastoma cell death through inhibition of an EGFR/AKT/SREBP-1/LDLR-dependent pathway. Cancer Discov. 1, 442–456 (2011).

52. T. Scully, N. Kase, E. J. Gallagher, D. LeRoith, Regulation of low-density lipoprotein receptor expression in triple negative breast cancer by EGFR-MAPK signaling. Sci. Rep. 11, 17927 (2021).

53. H. Ying, H. Zheng, K. Scott, R. Wiedemeyer, H. Yan, C. Lim, J. Huang, S. Dhakal, E. Ivanova, Y. Xiao, H. Zhang, J. Hu, J. M. Stommel, M. A. Lee, A.-J. Chen, J.-H. Paik, O. Segatto, C. Brennan, L. A. Elferink, Y. A. Wang, L. Chin, R. A. DePinho, Mig-6 controls EGFR trafficking and suppresses gliomagenesis. Proc. Natl. Acad. Sci. U. S. A. 107, 6912–6917 (2010).

54. K. Parag-Sharma, A. Leyme, V. DiGiacomo, A. Marivin, S. Broselid, M. Garcia-Marcos, Membrane Recruitment of the Non-receptor Protein GIV/Girdin (Gα-interacting, Vesicle-associated Protein/Girdin) Is Sufficient for Activating Heterotrimeric G Protein Signaling. J. Biol. Chem. 291, 27098–27111 (2016).

55. P. Ghosh, A. O. Beas, S. J. Bornheimer, M. Garcia-Marcos, E. P. Forry, C. Johannson, J. Ear, B. H. Jung, B. Cabrera, J. M. Carethers, M. G. Farquhar, A G{alpha}i-GIV molecular complex binds epidermal growth factor receptor and determines whether cells migrate or proliferate. Mol. Biol. Cell. 21, 2338–2354 (2010).

56. U. Dionne, A.-C. Gingras, Proximity-Dependent Biotinylation Approaches to Explore the Dynamic Compartmentalized Proteome. Front Mol Biosci. 9, 852911 (2022).

57. S. Saha, A. A. Anilkumar, S. Mayor, GPI-anchored protein organization and dynamics at the cell surface. J. Lipid Res. 57, 159–175 (2016).

58. Y. Luo, Y. Yang, P. Peng, J. Zhan, Z. Wang, Z. Zhu, Z. Zhang, L. Liu, W. Fang, L. Zhang, Cholesterol synthesis disruption combined with a molecule-targeted drug is a promising metabolic therapy for EGFR mutant non-small cell lung cancer. Transl Lung Cancer Res. 10, 128–142 (2021).

59. C. Chang, X. Tang, D. Mosallaei, M. Chen, D. T. Woodley, A. H. Schönthal, W. Li, LRP-1 receptor combines EGFR signalling and eHsp90α autocrine to support constitutive breast cancer cell motility in absence of blood supply. Sci. Rep. 12, 12006 (2022).

60. G. G. Hesketh, J.-Y. Youn, P. Samavarchi-Tehrani, B. Raught, A.-C. Gingras, Parallel Exploration of Interaction Space by BioID and Affinity Purification Coupled to Mass Spectrometry. Methods Mol. Biol. 1550, 115–136 (2017).

61. G. Liu, J. Zhang, B. Larsen, C. Stark, A. Breitkreutz, Z.-Y. Lin, B.-J. Breitkreutz, Y. Ding, K. Colwill, A. Pasculescu, T. Pawson, J. L. Wrana, A. I. Nesvizhskii, B. Raught, M. Tyers, A.-C. Gingras, ProHits: integrated software for mass spectrometry-based interaction proteomics. Nat. Biotechnol. 28, 1015–1017 (2010).

62. D. Kessner, M. Chambers, R. Burke, D. Agus, P. Mallick, ProteoWizard: open source software for rapid proteomics tools development. Bioinformatics. 24, 2534–2536 (2008).

63. D. N. Perkins, D. J. Pappin, D. M. Creasy, J. S. Cottrell, Probability-based protein identification by searching sequence databases using mass spectrometry data. Electrophoresis. 20, 3551–3567 (1999).

64. J. K. Eng, T. A. Jahan, M. R. Hoopmann, Comet: an open-source MS/MS sequence database search tool. Proteomics. 13, 22–24 (2013).

65. E. W. Deutsch, L. Mendoza, D. Shteynberg, T. Farrah, H. Lam, N. Tasman, Z. Sun, E. Nilsson, B. Pratt, B. Prazen, J. K. Eng, D. B. Martin, A. I. Nesvizhskii, R. Aebersold, A guided tour of the Trans-Proteomic Pipeline. Proteomics. 10, 1150–1159 (2010).

66. D. Shteynberg, E. W. Deutsch, H. Lam, J. K. Eng, Z. Sun, N. Tasman, L. Mendoza, R. L. Moritz, R. Aebersold, A. I. Nesvizhskii, Mol. Cell. Proteomics, in press.

67. V. Demichev, C. B. Messner, S. I. Vernardis, K. S. Lilley, M. Ralser, DIA-NN: neural networks and interference correction enable deep proteome coverage in high throughput. Nat. Methods. 17, 41–44 (2020).

68. J. D. R. Knight, H. Choi, G. D. Gupta, L. Pelletier, B. Raught, A. I. Nesvizhskii, A.-C. Gingras, ProHits-viz: a suite of web tools for visualizing interaction proteomics data. Nat. Methods. 14, 645–646 (2017).

69. J. Reimand, T. Arak, P. Adler, L. Kolberg, S. Reisberg, H. Peterson, J. Vilo, g:Profiler—a web server for functional interpretation of gene lists (2016 update). Nucleic Acids Res. 44, W83–W89 (2016).

